# High-Resolution Spatial Mapping of Electrocorticographic Activities with a 4096-Channel, Multiplexed Flexible Thin-Film Transistor Array

**DOI:** 10.1101/2024.08.18.608446

**Authors:** Yang Xie, Zepeng Zhang, Muyang Liu, Jinhong Guo, Miao Xu, Lei Zhou, Lei Zhang, Lihao Yao, Xiaolin Zhou, Zhengwei Hu, Liang Ma, Xiaojian Li, Yongxiang Guo, Jiaxin Lei, Yue Cao, Milin Zhang, Huachun Wang, He Ding, Xin Fu, Quanlei Liu, Yihe Wang, Lan Yin, Guoguang Zhao, Xing Sheng

**Affiliations:** Department of Electronic Engineering, Beijing National Research Center for Information Science and Technology, Institute for Precision Medicine, Laboratory of Flexible Electronics Technology, IDG/McGovern Institute for Brain Research, Tsinghua University, Beijing, 100084, China; Nanjing Research Institute of Electronics Technology, Nanjing, 210039, China; NeuCyber NeuroTech (Beijing) Co. Ltd, Beijing, 102206, China; The Institute of Polymer Optoelectronic Materials and Devices, South China University of Technology, Guangzhou, 510640, China; Guangzhou New Vision Optoelectronic Company Ltd., Guangzhou, 510530, China; School of Biomedical Engineering, Capital Medical University, Beijing, 100069, China; Chinese Institute for Brain Research, Beijing (CIBR), Beijing, 102206, China; Shenzhen-Hong Kong Institute of Brain Science-Shenzhen Fundamental Research Institutions, Shenzhen Institute of Advanced Technology, Chinese Academy of Sciences, Shenzhen, 518055, China; Shenzhen We-Linking Medical Technology Co., Ltd. Shenzhen, 518000, China; Department of Electronic Engineering, Tsinghua University, Beijing, 100084, China; Department of Electronic Engineering, Institute for Precision Medicine, Tsinghua University, Beijing, 100084, China; School of Integrated Circuits, Shenzhen Campus of Sun Yat-sen University, Shenzhen, 518107, China; Beijing Engineering Research Center of Mixed Reality and Advanced Display, School of Optics and Photonics, Beijing Institute of Technology, Beijing, 100081, China; School of Materials Science and Engineering, The Key Laboratory of Advanced Materials of Ministry of Education, State Key Laboratory of New Ceramics and Fine Processing, Laboratory of Flexible Electronics Technology, Tsinghua University, Beijing, 100084, China; Department of Neurosurgery, Xuanwu Hospital, Capital Medical University, Beijing, 100053, China

## Abstract

Advanced brain-machine interfaces (BMIs) demand implantable devices that offer flexibility, high-density and high-throughput capabilities. Conventional neural electrodes are constrained by the wiring strategy. Here we present a flexible and implantable electrocorticography (ECoG) device array (NeuroCam) based on metal-oxide semiconductor thin-film transistors (TFTs), to record brain activities at a large scale. Employing a multiplexing technique, the system is capable to record ECoG signals with up to 4096 channels and a density of 44 sites/mm^2^, while compressing the fan-in/fan-out leads to around a hundred. In a rabbit model with epilepsy, the NeuroCam array maps abnormal spike-wave discharges with an exceptional spatial resolution (150 μm) across extensive brain areas. The device strategy provides a promising route towards high-throughput BMIs with potential applications in fundamental neuroscience studies and practical biomedicine.

**Significance Statement:** Developing high-throughput neural recording techniques is critical for next-generation brain-machine interfaces (BMIs), but conventional neural electrodes are limited by the difficulty of scaling the lead wires. Here we report an implantable neural interface, based on a flexible thin-film transistor (TFT) array named NeuroCam. Employing an actively switching (multiplexing) technique, NeuroCam is capable to record electrocorticographic (ECoG) signals with up to 4096 channels and a density of 44 sites/mm^2^, with only 128 input/output leads. Validated in a rabbit model of epilepsy, NeuroCam records spatially resolved neural activities across extensive cortical areas with a resolution of 150 μm, and precisely maps the temporal evolution of epileptic waves. NeuroCam shows a promising solution for high-performance BMIs.

## INTRODUCTION

High-performance brain-machine interfaces (BMIs) represent powerful tools in fundamental neuroscience studies and promise broad applications in healthcare(1–3). In particular, realizing precise recording with large populations of neurons has been a long-standing goal in developing next-generation BMI systems(1, 2, 4, 5). Recently, emerging technologies in the materials design and device fabrication enable miniaturized wearable and implantable BMIs to record brain activities including electroencephalography (EEG)(6), electrocorticography (ECoG)(7–9), local field potential (LFP)(10), extracellular and intracellular spikes(11, 12), in non-invasive and minimally invasive manners. Specifically, flexible ECoG devices placed on the cerebral cortex provide a nonpenetrating means to measure population-level coordinated neural activities across wide cortical areas, and have been successfully applied for speech synthesis(13–15), motor decoding(16–18), and seizure localization(19). While conventional ECoG electrodes are typically a few millimeters in size, cortical columns sharing certain functional and anatomical properties have lateral dimensions ranging from tens to hundreds of micrometers(20). Moreover, experimental evidences prove that high-resolution ECoG recordings markedly improve the accuracy of neural decoding(21). Therefore, tremendous efforts focus on the innovation of thin-film, high-density microscale ECoG (μECoG) electrode arrays with over hundreds and even thousands of readout channels to obtain high-throughput brain activity mapping at a submillimeter resolution(9, 22). However, in passive neural electrodes, each recording channel requires at least one independent wire for signal readout, imposing formidable challenges for device fan-in/fan-out and scaling up high-density arrays(5, 22).

By contrast, active neural electrode arrays resolve the above-mentioned issue by time-division multiplexing, offering a viable solution to achieving high-density, high-throughput μECoG devices(23–25). Representative works include the development of flexible μECoG recording arrays integrated with thin-film transistor (TFT) based multiplexers made of monocrystalline silicon (Si), graphene and organic semiconductors(23, 25–29). Nevertheless, the production of these devices involves sophisticated or non-standard manufacturing processes, impeding their cost-effective fabrication with uniform performance on a large scale. Alternatively, metal oxide semiconductor based TFTs, which have been widely used in display technologies, are fully compatible with low-cost, meter-scale production on flexible platforms and provide a promising solution to large-area, multiplexed biological sensing(30). Prominent examples involve recently reported indium gallium zinc oxide (IGZO) TFT arrays for in vitro cultured cell sensing(31) or in vivo μECoG recording(32, 33); however, flexible high-throughput (> 1000 channels) μECoG devices made of oxide-based TFTs have not yet been realized for in vivo brain recording.

In this study, we report a flexible, multiplexed high-density μECoG array named NeuroCam for high-throughput brain activity mapping. NeuroCam features 4096 recording channels and an electrode density of 44 sites/mm^2^ based on a metal-oxide TFT array. The actively multiplexed scheme operates with only 128 input/output wires, effectively addressing the interconnection difficulties. In vitro and in vivo experiments demonstrate ideal biocompatibilities of the flexible device system. Validated in a rabbit model of epilepsy, NeuroCam records spatially resolved neural activities across extensive cortical areas. The high-density array precisely maps the temporal evolution of epileptiform spikes as well as the spatial distribution of the epileptogenic zone. This device concept provides a promising route towards high-density, high-throughput BMI systems compatible with industrial-scale production and brings unprecedented opportunities for both neuroscience and neuroengineering.

## RESULTS

### NeuroCam: A 4096-channel, multiplexed μECoG array

Presented in Figure 1, NeuroCam is based on a lanthanide-doped InZnO (Ln-IZO) TFT array(34) comprising 4096 pixel units arranged in a 64-row by 64-column grid, specifically designed for high spatial resolution ECoG acquisition. The manufacturing process is performed in an industrial-scale production line, which ensures desirable device scalability and uniformity over a large substrate (20 cm × 20 cm) with a high yield (Fig. S1). From bottom to top, the device structure (Fig. 1a) comprises a flexible polyimide (PI) substrate (25 μm thick), an active layer of Ln-IZO TFT array, multilayered organic/inorganic encapsulants and molybdenum (Mo) based interconnect electrodes, and gold (Au) based sensing pads. Fig. 1b shows schematic as well as cross-sectional scanning electron microscopic (SEM) images illustrating detailed device layouts. Among the TFTs with various channel dimensions, the device with a channel width/length of 200/10 μm exhibited the highest electrical conductance (Fig. S2). The fully fabricated flexible NeuroCam array has a total thickness of less than 30 μm and an active area of 9.6 mm × 9.6 mm (Fig. 1c). Each pixel measures 150 μm × 150 μm, and the size of the Au sensing pad is 70 μm × 135 μm (Fig. 1d and 1e). Overall, the NeuroCam array presents the highest channel count (4096) and density (44 sites/mm²) compared with state-of-the-art multiplexed μECoG techniques that have been validated in in vivo experiments (Fig. 1f and Table S1). Based on electrical characterizations in Fig. 1g and S3, we determine that the Ln-IZO TFT has a mobility of 15.13 cm^2^/V·s and a threshold voltage of 2.7 V. When turn-on voltage and turn-off gate voltages (*V*_gs_) are set at 6 V and −2 V, respectively, the TFT achieves an on/off ratio of about 6 × 10^6^. Additionally, transfer characteristic curves measured for multiple TFTs (*V*_ds_ = 0.1 V) manifest the device uniformity, and the channel impedance is determined to be 31.8 ± 6.4 kΩ at the turn-on state (Fig. 1h). The device’s frequency dependent impedance is further measured in the PBS solution, and its value is around 100 kΩ at 1–100 kHz (Fig. S3f).

**Figure 1.**
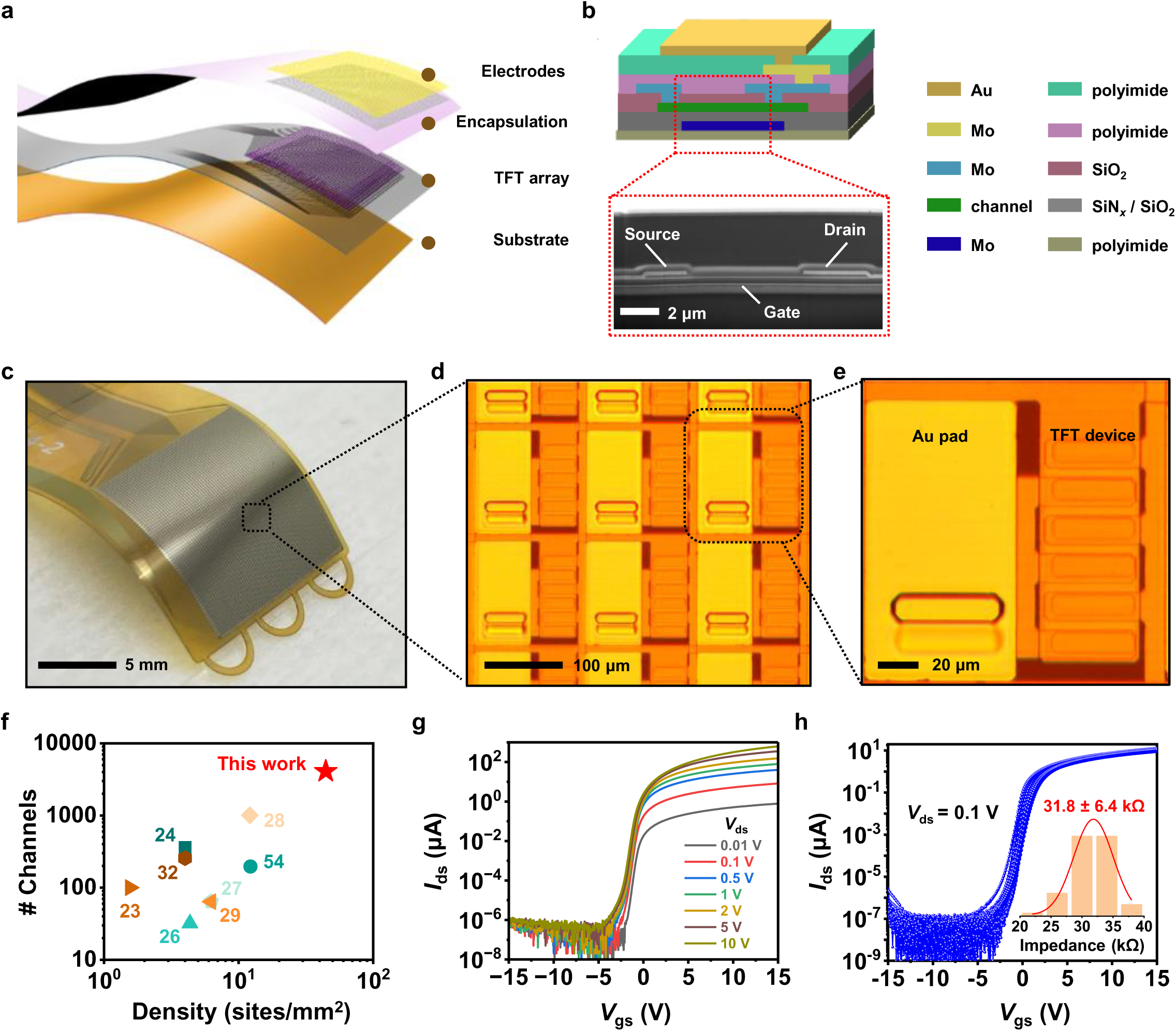
The NeuroCam: A flexible, 4096-channel multiplexed μECoG device array. **a.** Exploded view of the device structure, mainly comprising four layers: electrodes, encapsulation, TFT array, and substrate. **b.** Schematic diagram and corresponding cross-sectional SEM image showing the detailed structure of a single recording unit. The topmost Au pad serves as the sensing electrode, connected to the TFT drain electrode through a stepwise via structure made of multilayered Mo based interconnects and organic/inorganic encapsulation layers. The bottom substrate is made of PI (25 μm thick). **c–e.** Optical and zoomed-in microscopic photographs of a NeuroCam array. The dimensions of each pixel unit and corresponding Au sensing pad are 150 μm × 150 μm and 135 μm × 70 μm, respectively. **f.** Comparison of the NeuroCam array with existing multiplexed μECoG arrays used for in vivo experiments, in terms of their channel number and density. **g.** Transfer characteristic curve of a representative TFT under different *V*_ds_. **h.** Transfer characteristic curves of multiple TFTs in the array (*n* = 69 devices, *V*_ds_ = 0.1 V). Inset: Measured impedance of each device channel at a gate voltage *V*_gs_ = +6 V, with a statistical result of 31.8 ± 6.4 kΩ.

### In vitro characterization of the NeuroCam array

Figure 2 describes the operational scheme and in vitro performance of the NeuroCam multiplexed array. For a single TFT, the drain (D) port with the sensing electrode contacts with the targeted brain tissue, the source (S) port connects to the external output line, and the conduction between D and S ports is controlled by toggling the gate (G) port between turn-on and turn-off states (Fig. 2a). For the full multiplexed array, 64 S lines are connected to a multichannel recording system, with each line shared by 64 different channels sequentially scanned by corresponding G lines, allowing the system to operate with an overall 128 active input/output lines (Fig. 2b). Such a multiplexing strategy considerably mitigates the complexity of interconnects with external circuits. During the data acquisition, the specified G lines are sequentially set to the on state, activating the corresponding channels and cycling through all channels to complete time-division multiplexed signal recording (Fig. 2c).

**Figure 2.**
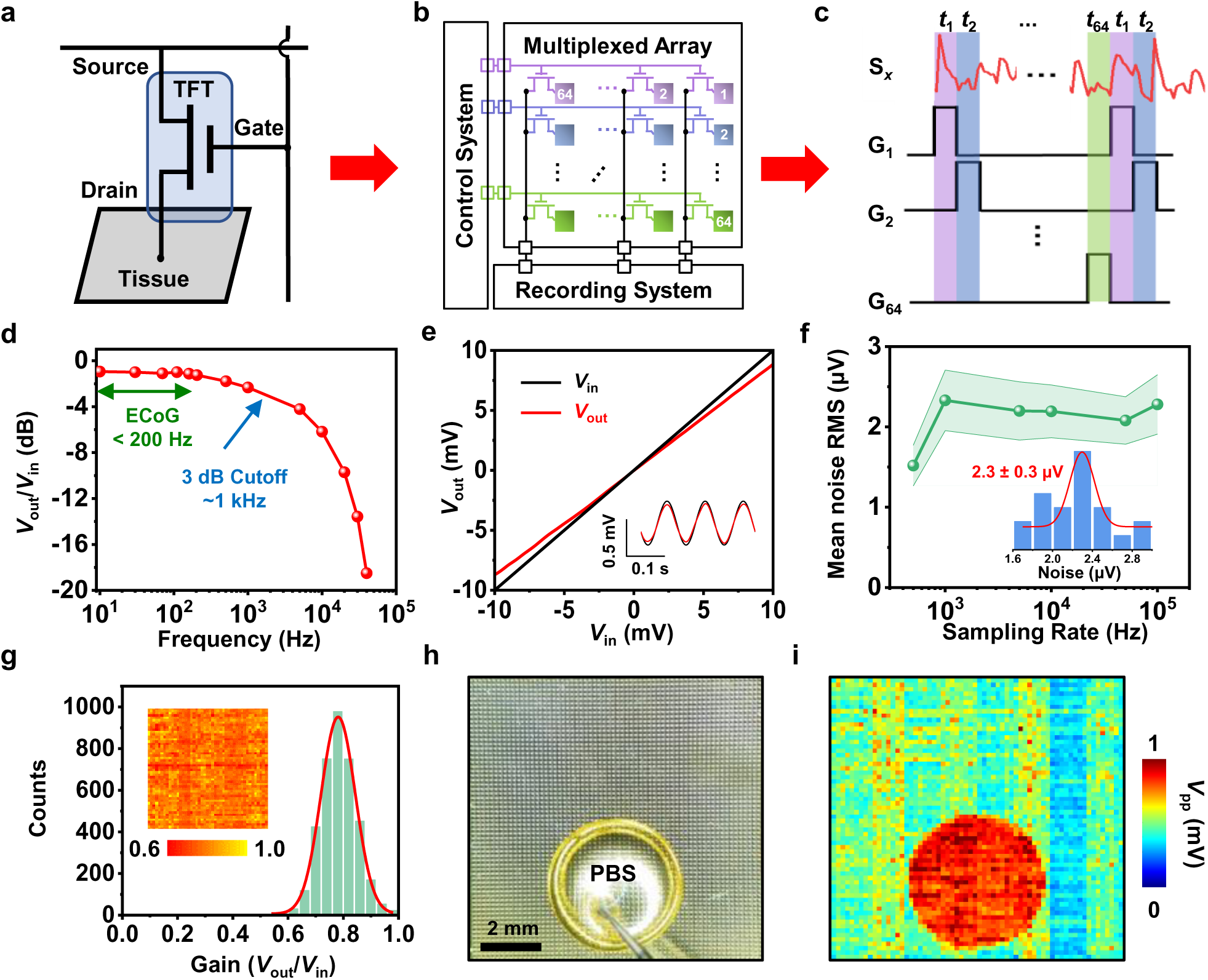
In-vitro characterization of the NeuroCam array. **a–c.** Operational principle of the NeuroCam array. **a.** Single unit: The drain electrode of the TFT device directly contacts with brain tissue, the gate controls the conduction between the drain and the source, and the source connects to the external recording system. **b.** TFT array: The recording system continuously collects data from all the source (S) lines, while the control system modulates all the gate (G) lines, by sequentially turning on one G line (*V*_on_ = +6 V) and keeping other G lines at the off state (*V*_off_ = −2 V). **c.** Collected results: The time-domain control signals (G_1_ to G_64_) segment the collected data, enabling time-division multiplexed recording in a specific S line (S*_x_*) cyclically repeating from *t*_1_ to *t*_64_ (*t* = 62.5 μs for each channel). **d.** Measured frequency characteristics of a recording channel, showing a 3 dB cutoff at ∼1 kHz. Here the G line is kept on, while a 10 mVpp sinusoidal wave signal of varying frequencies is applied to the drain exposed in PBS. **e.** Measured relation between the input and output peak signals (*V*_out_ vs. *V*_in_). Inset: corresponding signal waves of *V*_out_ and *V*_in_ (0.6 mVpp). **f.** Statistical results of mean root-mean-square (RMS) noise for 23 active channels in the array. Inset: Histogram and Gaussian fit curve of noise levels for these channels at a sample rate of 100 kS/s. **g.** Gain (*V*_out_/*V*_in_) characteristics of all the 4096 channels in the array in PBS. The input signal is a sine wave with an amplitude of 1 mV and a frequency of 12 Hz. **h.** Photograph of a NeuroCam array with a droplet of PBS on its surface. **i.** Mapping results of spatially resolved output signals when injecting a 1 mVpp sine signal (18 Hz) into PBS.

We characterize in vitro performance of NeuroCam by immersing the array in the phosphate buffered saline solution (PBS), where a platinum (Pt) based mesh electrode supplied input electrical signals with varied frequencies and amplitudes (Fig. S4). Encapsulations protect the full device array from the solution, except the Au sensing pads that connect with D ports of TFTs. Fig. 2d plots the frequency response of *V*_out_/*V*_in_ for a typical TFT in the array, which is collected by changing the input signal (*V*_in_) frequency from 10 Hz to 40 kHz and measuring the collected output voltage (*V*_out_) via the S port (Fig. 2e). The obtained 3 dB attenuation frequency exceeds 1 kHz, which is much higher than the typical frequency range of ECoG signals (< 200 Hz). We further evaluate the device’s noise levels in both static and dynamic modes. In the static mode, one G line remains open while others are closed, and noise levels are determined at different sampling rates and precision modes (Fig. S5). When operated in the high-precision mode, the recording system is capable to collect data from a maximum of 24 S lines. In this case, the system reaches the lowest root mean square (RMS) noise level of 2.3 ± 0.3 μV at a sampling rate of 100 kS/s (Fig. 2f). Similar to previous works(35), this static mode is specifically designed for high-precision recordings from small regions within the array’s coverage and demonstrates the lowest achievable noise level under controlled conditions. It provides a reference baseline for noise performance, allowing the adjustment of active channels depending on the application. Dynamic scanning performed in the full 64 × 64 array leads to elevated noise levels (Fig. S6) comparable to previously reported μECoG arrays with similar multiplexing systems(24, 28, 36). While our system records a greater number of channels (4096) than those in prior works (typically 256 or 1000 channels), its noise level (63.6±24.1 µV_rms_) is within an acceptable range when considering the number of channels. When reducing the number of recording channels, the noise level becomes comparable to that of other active ECoG devices. For example, reducing the channels to 384 (24×16) makes the system noise to be 8.8±4.2 µV_rms_., similar to a previously reported 256-channel array (2.65 µV_rms_) (32). When operating 1536 channels (24×64), the noise is 12.6±8.4 µV_rms_, which is lower than a previously reported 1008-channel array (58 µV_rms_) (28).

In such a multiplexed array, each 64 input and 64 output channels share a same gate line and a same source line, respectively. Therefore, crosstalk signals may exist among pixels. In Fig. S7 and Movie S1, we evaluate the system’s crosstalk by applying an input signal on a single pixel and mapping the entire array. Adjacent pixels in the direction of source line receives higher crosstalk signals (−6.3 dB) comparing to those in the direction of gate line (−18.6 dB), while second nearest neighboring and further pixels show minimal crosstalk signals (−22.5 dB).

The histogram of NeuroCam illustrates uniformly distributed performance among 4096 channels, with a gain (*V*_out_/*V*_in_) of 0.78 ± 0.12 (Fig. 2g). Additionally, we emulate spatially resolved bioelectric signals by applying a sinusoidal wave (1 mVpp, 18 Hz) via a PBS droplet on the array (Fig. 2h). The mapping results clearly demonstrate the NeuroCam’s capability of dynamically recording electric signals with desirable spatial and temporal resolutions in the aqueous environment (Fig. 2i, Fig. S8 and Movie S2). In the absence of PBS, baseline similarities appear in some pixels due to the backend acquisition system, which uses 8 ADCs with 4 channels each, leading to similar baseline signals in channels with matching numbers across different ADCs. Other complex shape of PBS droplets can be formed and recorded with the NeuroCam, with an example shown in Fig. S9 and Movie S3.

Furthermore, we conduct systematic characterizations to evaluate the NeuroCam’s biocompatiblites. First, cytotoxicity tests show no significant difference in cell viability between the TFT array and glass (Fig. S10). Second, the infrared thermal imaging reveals negligible temperature rises upon the device operation, suggesting that the device array would not impose any adverse thermal effects on the brain tissue (Fig. S11). Third, the flexible NeuroCam array maintains stable performance under deformation, with device yield and gain changed slightly after 10,000 bending cycles (Fig. S12). Last, the array presents modest chronic reliability, with the device yield changing from 100% to ∼80% and a stable gain among all the channels after soaking in PBS solution at 37 °C for two weeks (Fig. S13).

### High-throughput, in vivo mapping of epileptiform activities using NeuroCam

Epilepsy is a common neurological disorder marked by abnormal neuronal discharges in the brain(37–39). Monitoring these neural activities across wide brain regions with high spatial resolution is crucial for understanding mechanisms of epileptogenesis, identify the epileptogenic zone, and improving treatment strategies(9, 19, 24, 40). Here we utilize the NeuroCam array to record ECoG signals in a rabbit model with epilepsy (Fig. 3a). In anesthetized animals, intracortical administration of penicillin induces focal epilepsy, and the subdurally positioned 4096-channel NeuroCam array covers a wide cortical area including the injection site (Fig. 3b and Fig. S14). Figs. 3c and 3d present ECoG signals as well as corresponding and their time-frequency patterns registered by a representative channel near the injection site at different time points. Fig. S15 shows the noise characteristics of the NeuroCam system before the penicillin injection. It should be noted that these signals encompass the noise associated with the device array and the recording system, as well as the animal’s ECoG activity at the resting state. These recorded epileptic activities are in accordance with previous reports of intracortical penicillin administration in rodents(10, 41, 42). Following a brief quiescent period (∼3 minutes) post-penicillin administration, epileptic seizure activities start and gradually intensify, and eventually reach status epilepticus that could last for more than 20 minutes. The status epilepticus is characterized by typical spike-and-wave discharges with a frequency of 1–4 Hz (due to individual differences in the epileptic model). Fig. 3e plots the evolution of spike frequency after injection for different experiment trails in four animals. All four animals exhibit increased spike rates after injection, although their values (1–4 Hz) are different owing to the variations of physiological and environmental states. These rhythmic spike patterns could be crucial indicators of underlying brain tissue lesions(43, 44).

**Figure 3.**
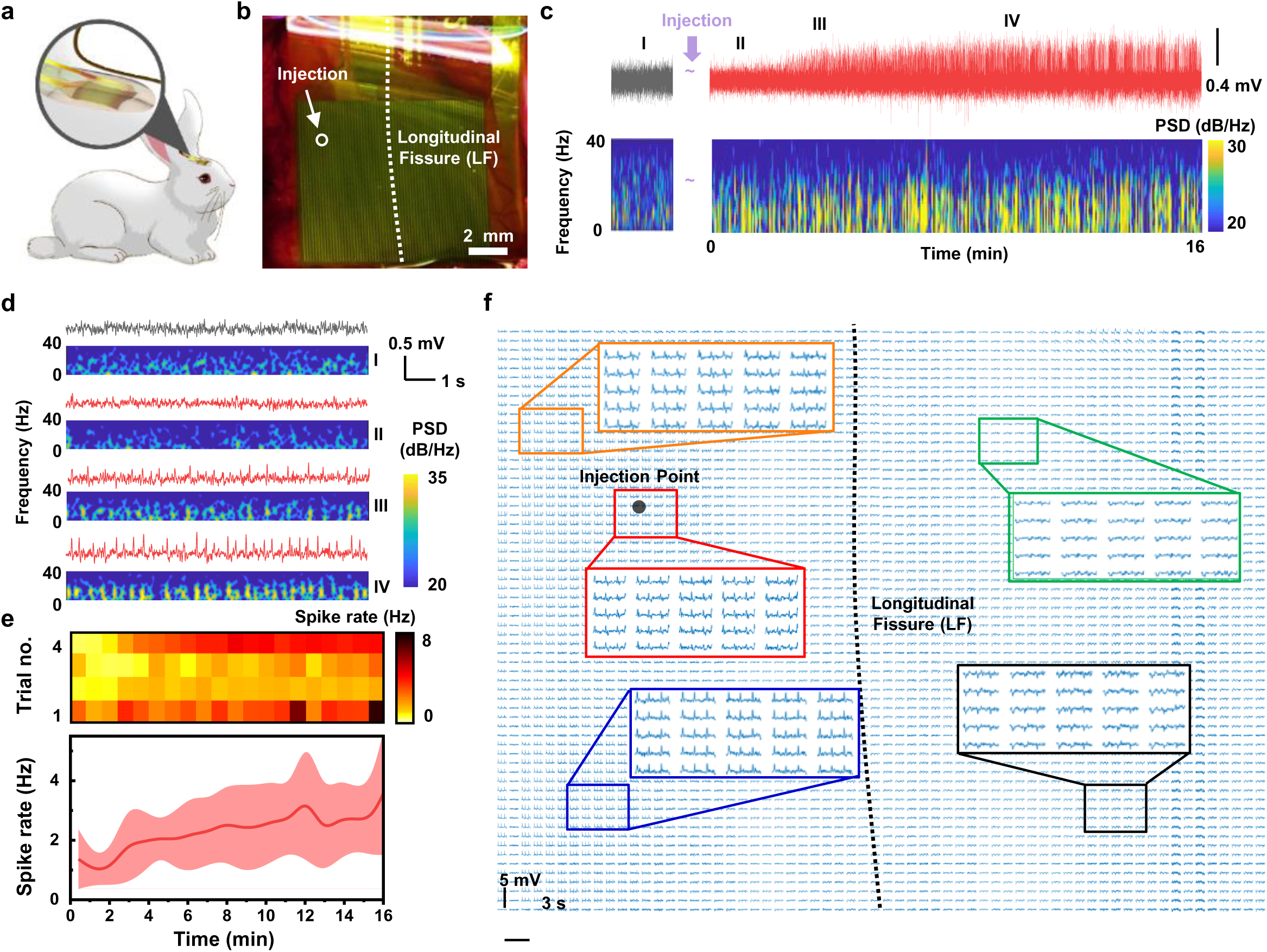
In vivo recording of epileptiform activity in rabbits using the NeuroCam array. **a.** Schematic diagram and **b.** top-view photograph illustrating a NeuroCam array mounted on the cortex of a rabbit. The arrow indicates the location of penicillin injection. **c.** Typical ECoG signals and their time-frequency patterns registered by a representative channel near the injection site before and during epileptic seizures. **d.** Temporally resolved ECoG signals in c. and corresponding time-frequency patterns at four different time points: (I) before penicillin injection, (II) shortly after injection, (III) onset of epilepsy, and (IV) status epilepticus. **e.** Dynamic response of epileptic spike-wave discharges: heatmap of four individual trials from *n* = 4 rabbits (top); averaged spike rates (bottom). **f.** ECoG activities recorded from all 4096 channels in the array, during status epilepticus in the rabbit. Insets show enlarged plots from different brain regions.

The full mapping results from all the 4096 channels in the array, as shown in Figure 3f and Figure S16, clearly illustrate spatial differentiation of ECoG activities. Electrodes near the injection site exhibit similar activity, indicating the propagation of synchronized epileptic discharges, which may also be influenced by crosstalk from the multiplexed scanning system. Among the various electrophysiological characteristics of epileptic seizures, focal intense abnormal activity is the most specific biomarker(44). The regions with most intense spike signals closely correlate with the location of the injection point. In these active regions, bilateral spikes occur with a burst frequency of ∼3 Hz, a duration of ∼150 ms, and a peak-to-peak amplitude of ∼2 mV. By contrast, ECoG signals collected by channels distant from the injection point (specifically, in the contralateral hemisphere) exhibit much weaker activities.

### Analysis of epileptic seizures based on high-throughput ECoG mapping

The NeuroCam array provides a large coverage and high spatial resolution ECoG mapping, enabling the capture of intricate local and global dynamics during epileptic seizures on the cortical surface. This reflects the characteristic large-scale synchronized discharges during epileptic activity. These recorded epileptiform discharges feature spike waves with a high amplitude (∼2 mV) and a high signal-to-noise ratio (SNR = 10 ×log_10_ (Signal Power / Noise Power) ∼ 25 dB) (Fig. 4a and 4b). The high-density mapping across extensive brain regions reveals detailed spatial and temporal evolution of seizure activities (Fig. 4c, S17, S18, Movie S4). In epileptic seizures, ECoG signal propagation is accompanied by large-scale synchronized discharges, with regions often spanning millimeter scales (45). Despite the array’s 150-micron pitch, the larger spatial scale of the epileptic focus likely results in observed activity islands near 1 millimeter. However, detailed analysis of the mapping images reveals clear submillimeter differences between channels. These time-lapse mapping results clearly illustrate the developmental process of abnormal electrical discharges during an epileptic event across the brain: (i) The seizure spike initially arises near the focal area; (ii) The focal area shows a positive potential that propagates abnormal signals outward, with the positive signal spreading to the lower-left ipsilateral brain region and the negative potential extending to the contralateral hemisphere; (iii) Abnormal electrical waves spread throughout the entire area covered by the array, with the focal area shifting to a negative signal and gradually spreading the same waveform to adjacent ipsilateral regions; (iv) The electrical activity subsides, with only mild oscillations in certain regions of both hemispheres. This pattern of large spikes followed by slower waves of opposite polarity aligns with previously reported results of focal epilepsy(46, 47). Furthermore, the high-density and large-area ECoG mapping of NeuroCam reveals more extensive spatial information not commonly observed in earlier studies. Filtering the epileptic waveforms into different frequency bands exhibits distinct time-frequency oscillations associated with spike bursts (Fig. S19). Fig. 4d presents the RMS amplitude of a typical spike filtered through various frequency windows. The progression of spike waves is most prominent in the beta-band (12–30 Hz), clearly showing the origins and convergence areas of the abnormal ECoG activities. Additionally, owing to its ultrahigh channel density (44 sites/mm^2^), the NeuroCam array can achieve sub-millimeter precision in localizing epileptic foci by mapping the onset of seizure spikes (Fig. 4e). Based on the ECoG recording, we can extract both spatially distributed RMS amplitudes and signal delays collected from three different rabbits (Fig. S20). By comparing these two different maps, we observe that the seizure onset zone aligns more closely with the penicillin injection site rather than the region with the highest spike amplitude. Notably, the color bar in the heat map is specifically chosen to clearly show the active regions, which enhances the visibility of the signal dynamics during the epileptic event.

**Figure 4.**
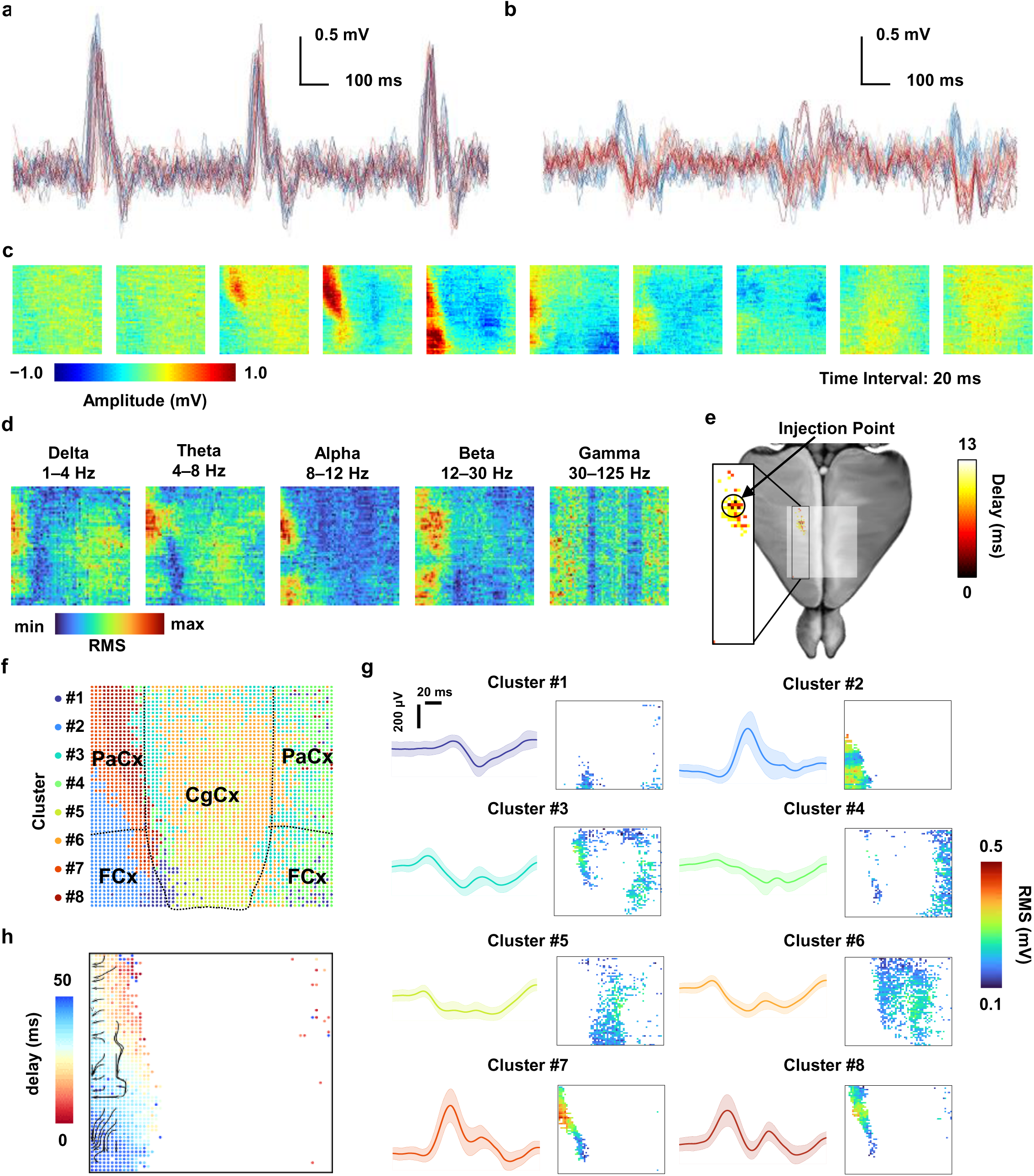
High-spatial-resolution ECoG mapping of rabbit cortex under epileptic seizures. **a, b.** Recorded waveforms for (a) channels adjacent to the injection site and (b) channels distant from the location in separate diagrams. These recurring rhythmic spikes (2–4 Hz frequency) indicate ongoing epileptic activity. **c.** Time-lapse ECoG mapping of an individual spike shown in (b), with a total duration of 180 ms. **d.** RMS heatmaps of waveforms in different frequency bands. **e.** Alignment of the delay mapping with device positions enables sub-millimeter precision localization of the epileptic focus (dorsal view). **f.** Clustering results of waveforms from all active channels in the array showing the spatial distribution of 8 clusters with distinct waveforms, overlaid with segmented cortical areas of the rabbit brain (dorsal view). **g.** Waveforms (left) and corresponding RMS heatmaps (right) of the 8 delineated clusters. **h.** Heatmaps indicating peak delays of clusters 2, 7, and 8, associated with abnormal high-amplitude spikes related to induced epileptic activities. The streamline plots illustrate propagating waves, with vectors representing the direction of wave propagation.

The brain-wide mapping also helps classify ECoG signals of different characteristics in different brain regions. Methods based on principal component analysis (PCA) and k-means clustering categorize the signals registered from all 4096 channels into 8 clusters, which are mapped at their respective locations (Fig. 4f). These 8 distinct clusters feature various spike waveforms such as biphasic spikes and spike-and-slow wave complexes (Fig. 4g), which exhibit strong spatial correlation with segmented cortical areas of the rabbit brain(48). In particular, channels showing intense spike activities (clusters 2, 7, and 8) are mostly located in the ipsilateral hemisphere of the lesion, primarily the parietal cortex (PaCx) and the frontal cortex (FCx). These clusters distinctly delineated the boundary of the cingulate cortex (CgCx). Finally, the NeuroCam array also captures detailed spatiotemporal development of epileptiform discharge waves at the submillimeter scale, depicting the progression of brain waves spreading outward from the focal area (Fig. 4h).

ECoG signals in the gamma band with frequencies >30 Hz are associated with advanced cognitive functions such as attention, memory, and perception, and commonly used to infer underlying neural processing mechanisms(49, 50). Additionally, patterns of gamma oscillations (30–70 Hz) can be utilized to locate epileptogenic regions in focal cortical dysplasia (51). Figure 5 depicts a 5-second segment of full-band raw data (1–125 Hz) and gamma (30–125 Hz) waveforms recorded during penicillin induced status epilepticus. Continual spike bursts in the raw data indicate an ongoing epileptic seizure (Fig. 5a). Considering a typical epileptic spike (duration 20 ms), mapping results of both raw data and gamma rhythms clearly illustrate alternating positive and negative waves originating from the epileptic focus and spreading to surrounding areas (Fig. 5a, 5b). Furthermore, RMS mapping results of these waveforms highlight the most active area and indicate distinct spatial coverage related to the epileptic focus area (Fig. 5c, 5d). Specifically, the RMS mapping of the gamma band signal locates an active area closer to the injection site when compared to the raw data, underscoring the effectiveness of gamma signals in pinpointing epileptic foci. Although gamma signals generally have lower amplitudes than other low-frequency ones, they can still effectively localize the epileptic focus, as shown by the mapping results.

**Figure 5.**
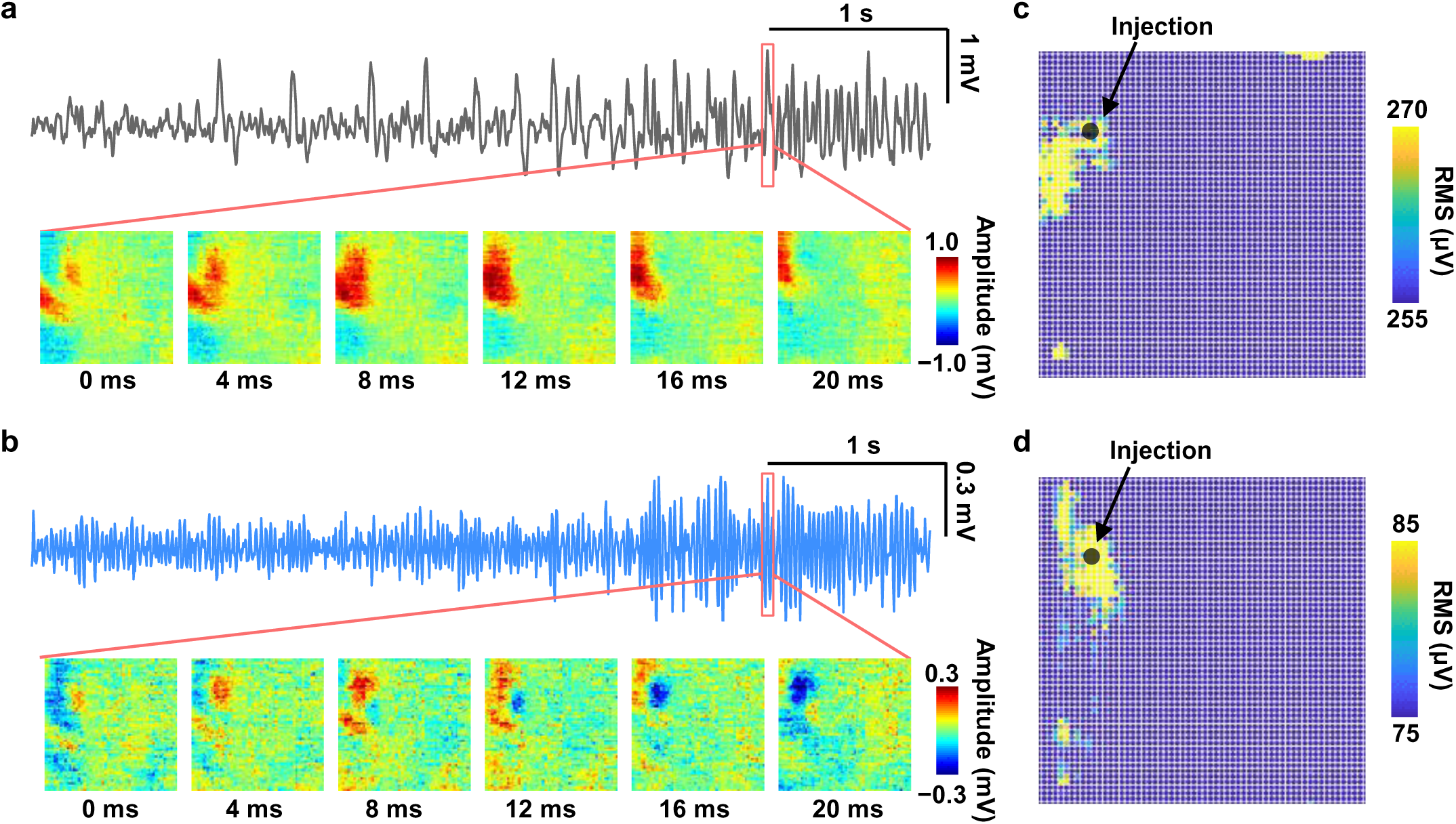
NeuroCam records gamma band signals that more precisely locate the epileptic focus. **a, b.** Raw data (a) and gamma band (b) waveforms registered by a representative channel near the injection site over a 5-second period, with time-lapse mapping results during an epileptic spike (duration 20 ms, interval 4 ms). **c, d.** RMS mapping results of the raw data waveform (c) and the gamma band signal (d). The arrow indicates the location of penicillin injection. The range of color bars are chosen to make the seizure-prone area more distinct.

We also assess the spatial correlation of ECoG signals collected from all the 4096 channels in the NeuroCam array (Fig. S21). Specifically, we analyze the correlation coefficients at varied channel distances for the 5-second epilepsy signals in Fig. 5. Generally, the correlation decreases as the inter-channel distance increases, consistent with previous studies(21, 52). Notably, the high-density NeuroCam array allows for spatial analysis down to the sub-millimeter level (as low as 150 μm). Correlation coefficients detected by channels in the G and S directions have similar dependence on the inter-channel distance (Fig. S21a). Averaged correlation coefficients between channels spaced more than 300 μm apart in both G and S directions are below 0.7, suggesting that the high-density array is meaningful for precise ECoG recording. For distances greater than 1.2 mm in the G direction and 1.8 mm in the S direction, the coefficients are below 0.5. The correlation among electrodes in the same S line is stronger than that in the same G line, indicating more crosstalk among channels on the same S line under multiplexed operation, in agreement with results in Fig. S7. Furthermore, ECoG signals in different frequency bands present different spatial correlations (Fig. S21b and S21c). As distance decreases, results of low-frequency signals quickly converge at the submillimeter scale, while the correlation of high-frequency signals in the gamma band is significantly weaker than those in other low-frequency bands (below 0.4 at 150 μm), suggesting that gamma signals are more spatially localized than other low-frequency signals. These characterizations further suggest that the high-density array with a submillimeter resolution would be more critical in recording high-frequency gamma waves.

## DISCUSSION

In this study, we develop the 4096-channel NeuroCam system based on flexible, active TFT arrays, and demonstrate its capability for high-throughtput ECoG recording across large brain areas in epileptic rabbits. Compared to previously reported multiplexed μECoG arrays demonstrated for in vivo studies, our NeuroCam system presents superior features in terms of its channel number (4096) and density (44 sites/mm^2^). Its high spatial resolution and extensive coverage provide practical advantages for precisely analyzing the spatiotemporal progression of epileptiform abnormalities and localizing epileptogenic zones. Additionally, understanding the differential and correlational patterns across brain regions during specific physiological activities can offer crucial insights in unraveling complex neural circuitry. On the other hand, the high-density, submillimeter electrodes would be particularly valuable in exploring correlations between cortical columns (in size from tens to hundreds of micrometers) and decoding complex behaviors like movement, sensing and vision(20, 43).

Since the multiplexing strategy have effectively resolved the wiring issue of passive μECoG electrodes, the mature manufacturing process of flexible TFTs could readily boost the channel number from the kilo scale (10^3^) to the mega scale (10^6^). Future improvements also involve optimizing the system’s signal-to-noise ratio, chronic stability and portability. Although the current array is able to map epileptic seizures, its noise level limits the precise detection of more intricate neural activities, such as sensory perception and high-order cognitive functions. Implementing sensing electrodes with lower impedance(53) and preamplifiers in the TFT arrays(24, 54, 55) would further decrease the system’s noise level down to several microvolts. Thinner and more flexible substrates could promote a more conformal attachment over the curved cortex(56), and advanced encapsulation and adhesive coatings(57, 58) could enhance the array’s long-term operation in living animals spanning weeks or even months. Customized ASIC chips, wireless power harvesting and data transfer schemes(59, 60) would further miniaturize the acquisition system and facilitate its portable use in freely moving scenarios. To summarize, the presented device design and production strategies pave the way for next-generation, high-performance BMI technologies and will bring unprecedented opportunities to neuroscience and neuroengineering.

## METHODS

### Device Fabrication

The device fabrication process was described as follows (Fig. S1): (1) A 200 nm thick Mo layer was sputtered onto a pre-cleaned PI substrate (25 μm thick) to form the bottom gate, followed by lithographic patterning; (2) Silicon nitride (SiN*_x_*) and silicon dioxide (SiO_2_) (200 nm / 200 nm) films were grown via plasma enhanced chemical vapor deposition (PECVD) as the gate insulating layers; (3) A 30 nm thick Ln-IZO semiconductor layer was sputtered as the TFT channel; (4) A 200 nm thick SiO_2_ layer was deposited via PECVD as the source/drain insulating layer; (5) An etch barrier layer was dry-etched and patterned to protect the active layer, followed by sputtered Mo films (200 nm thick) to form the source/drain electrodes; (6) A 2 μm thick PI layer was spin-coated and patterned as the organic insulating layer; (7) A 200 nm thick Mo film was sputtered and patterned to form interconnects; (8) A 2 μm thick PI film was spin-coated and patterned as the insulation; (9) 5 nm Ti/100 nm Au films was evaporated onto the drain electrode to form sensing pads; (10) The sample was cut into individual arrays using a laser scriber; (11) Flexible device arrays are peeled off from the glass substrate.

Subsequently, the array was connected to a flexible printed circuit (FPC) via anisotropic conductive films (ACFs) through thermal pressing. The other end of the FPC was inserted into a zero-insertion force (ZIF) connector on a printed circuit board (PCB) to link with the data acquisition (DAQ) system.

### Data Acquisition System

The DAQ system was configured using PXIe-6739 and PXIe-4309 modules (National Instruments Corp.), with the PXIe-6739 as the input module and the PXIe-4309 as the output module. The DAQ system connects to the flexible array (attached on the PCB) via very high-density cable interconnect (VHDCI) cables. The input and output modules were managed via a LabVIEW software program (National Instruments Corp.) developed to support multiple channel configurations (e.g. 64 ×64 = 4096, 23 ×64 = 1472, etc.) and physical sampling rates (e.g. 250 S/s, 1000 S/s, etc.). During experiments, the PXIe-4309 was set to a maximum single-channel sampling rate of 100 kS/s with an oversampling ratio of 6.25 to achieve higher precision. For scanning, a control pathway from the PXIe-6739 output was connected to the G1 line signal to provide a marker for multiplexed signal alignment, ensuring accurate signal synchronization. The entire system utilized three PXIe-4309 cards and one PXIe-6739 card. Each PXIe-4309 card contains eight ADCs, with each ADC supporting four acquisition channels. Typically, three PXIe-4309 cards were used to record signals from 64 S lines. For the higher precision mode, each PXIe-4309 could be set to acquire from no more than eight S lines, enabling the use of only one channel per ADC.

### In Vitro Characterization

#### Morphological Characterization

Optical images at various magnifications were captured using an optical microscope (XZJ-L2030, Phenix). Cross-sectional images of devices were obtained using FIB-SEM (QUANTA 200 FEG, FEI).

#### Electrical Characteristics of Individual TFTs

Transfer characteristics (*I*_ds_–*V*_gs_), output characteristics (*I*_ds_–*V*_ds_), and capacitance characteristics (*C*–*V*_gs_) of individual TFTs were measured using a semiconductor device parameter analyzer (Keysight B1500A, Keysight Technologies). The frequency response curve was obtained by applying 1 mVpp sine waves with different frequencies to the drain electrode and measuring *V*_out_/*V*_in_. Additionally, the TFT array was immersed in a PBS solution, and linear voltage (from −10 mV to +10 mV) or sinusoidal signals (500 μVpp, 12 Hz) were applied using a platinum mesh electrode. In vitro impedance (vs. frequency) of the TFT was measured in PBS with an electrochemical workstation (CHI 650E, Shanghai Chenhua Co., Ltd, China).

#### Biocompatibility Characterization

For the cell viability test, a small TFT array (1.6 × 1.6 mm^2^) was transferred on glass. Before cell culture, the sample was coated with 0.01% poly-l-lysine (No. P8920, Sigma-Aldrich) overnight, then coated with 1-2 μg cm^−2^ laminin (No. L2020, Sigma-Aldrich) for more than 4 hours. Dorsal root ganglia (DRG) of male rats (4 weeks) were cxcised and transferred in DMEM/F12 (No.11330-032, Gibco) on ice. Tissues were cut into small pieces and digested in an enzyme solution containing 5 mg ml^−1^ dispase (No.ST2339, Beyotime) and 1 mg ml^−1^ collagenase (No.ST2294, Beyotime) at 37 °C for 1 h. Then the samples were triturated with 1 ml pipette for 30 times until the liquid becoming turbid. After centrifugation at 250 g for 5 min, cells were washed in 2 ml of DMEM/F12 + 15% (w/v) bovine serum albumin (No.SRE0096-10G, Sigma) and resuspended in 2 ml of DMEM/F12 + 10% fetal bovine serum (No.F802A, i-presci scientific) + 1% penicillin–streptomycin (No.15140-122, Gibco). 10% of the total DRG cells were seeded on the TFT device or standard glass coverslip in each well of the 96 wells plate. The DRG cells were cultured in an incubator at 37 °C for 1 day. For the assay, the samples were labeled with Calcein/PI Cell Viability/Cytotoxicity Assay Kit (Beyotime). Live cells were labeled with Calcein AM (green), and dead cells were labeled with Propidium Iodide (red). After adding the dyes, the samples were incubated for 30 min at 37 °C with 5% CO_2_. The samples were washed with HBSS for three times. The viability (%) was calculated as total No. of live cells / total No. of cells ×100.

#### Mechanical Testing

The flexible array was mounted on an in-situ mechanical testing platform (IBTC-300SL, Care Corp.) for bending tests, with a bending radius set to 5 mm and repeated 10,000 times. The array was then immersed in a PBS solution and driven by a platinum mesh electrode with a 2 mVpp, 12 Hz sine wave. Using the DAQ system, the array was scanned in time-division multiplexing mode with 4096 channels activated at an equivalent sampling rate of 250 S/s. The number of undamaged channels and the average gain were recorded.

#### Noise Characterization

The array was immersed in 1× PBS solution (pH = 7.4, Sigma-Aldrich), with a platinum mesh electrode providing a ground signal. In the static mode, the DAQ system makes one G line active while keep other G lines closed, to record noise levels at different sampling rates. When recording from only 23 S lines one of the 24 available channels is reserved for capturing the switch calibration signal, and each of the remaining 23 ADCs was configured with a single acquisition channel to achieve higher precision and lower noise. When recording from 64 S lines, all 24 ADCs were used, with each ADC handling 2 or 3 acquisition channels. In the dynamic mode, noise measurements were conducted under three different configurations: 64 (S) × 64 (G) = 4096 channels at 250 S/s, 24 (S) ×64 (G) = 1536 channels at 250 S/s, and 24 (S) ×16 (G) = 384 channels at 1000 S/s. In each configuration, the total sampling rate for each S line was fixed at 100 kS/s.

#### In-vitro mapping testing

A drop of 1× PBS solution (pH = 7.4, Sigma-Aldrich) was placed on the array, and a 1 mVpp, 18 Hz sine wave was input using a tungsten probe. The DAQ system recorded data from the 4096-channel array at a sampling rate of 250 S/s.

### In vivo experiments

#### Animal Surgery

All animal procedures described in this study were approved by the Tsinghua University Institutional Animal Care and Use Committee (IACUC). Adult male New Zealand white rabbits (*Oryctolagus cuniculus*) (2000–2500 g) were purchased from Beijing Fangyuan Experimental Animal Technology Co., Ltd. and were housed individually under standard conditions. The animals were maintained on a 12-hour light/dark cycle at 22–25 °C, with ad libitum access to food and water.

After weighing the rabbits, they were anesthetized using a combination of Xylazine (6.5 mg/kg) and Zoletil (13 mg/kg) administered intramuscularly. Additional doses, at one-quarter of the initial dose, were given every 30 minutes during the procedure. Anesthesia was verified by pinching the animal’s hind limbs before starting the surgery. Meloxicam (0.3 mg/kg) was administered pre-operatively for pain relief, and erythromycin ointment or medical petroleum jelly was applied to protect the eyes. The anesthetized rabbit was secured in a stereotaxic frame and maintained on a warming pad to stabilize body temperature. The head was shaved, disinfected with 75% ethanol (Sigma-Aldrich), and the scalp was incised to expose the skull. The skull was aligned parallel to the reference plane of the positioning instrument, adjusted based on the rabbit’s cranial sutures. The exposed skull was etched with 8% hydrogen peroxide (Sigma-Aldrich). A craniotome with a 1.2 mm diameter was used to create a hole in the target area of the skull, with continuous saline irrigation to prevent overheating of the brain. A 1.4 mm diameter stainless steel screw was inserted as a reference electrode. The dura mater was removed using curved surgical forceps and scissors, and a sterile, saline-soaked cotton ball was placed over the exposed brain to maintain moisture.

#### Epilepsy Induction and Acute ECoG Recording

The skull screw was grounded with a metal wire and connected to the DAQ system. A glass syringe needle was implanted near the hippocampus (AP: 4 mm, ML: 2.5 mm, DV: −2.3 mm). Penicillin G potassium (1000 IU in 2 μL/min, MCE) was injected into the hippocampus at a concentration of 200 IU/μL. After removing the needle, the NeuroCam array were placed on the cortex and held with hemostatic sponges in place.

#### Array Characterization and Data Processing

All data processing and analysis were conducted using Origin (OriginLab) and MATLAB (MathWorks). Recorded data were initially segmented in MATLAB. The timing of array row gating was determined based on the marker signal. The average of the more stable 3rd to 5th data points from 6 or 7 points in each row was used as the sampling value. This process was repeated for each column, converting the recorded 64 columns of data into a 64 × 64 array. The data were then filtered and smoothed in Origin: a 49–51 Hz band-stop filter was used to remove power-line noise, followed by a 1–199 Hz band-pass filter to isolate the ECoG signals, and Savitzky-Golay smoothing was applied. Time-frequency plots, heatmaps, correlation analyses, and RMS analyses were performed using MATLAB functions such as ‘spectrogram’, ‘heatmap’, ‘corrcoef’, and ‘rms’. Gaussian fitting for histograms was done using Origin’s Gaussian fitting function. Signals were divided into Delta (1–4 Hz), Theta (4–8 Hz), Alpha (8–12 Hz), Beta (12–30 Hz), and Gamma (30–125 Hz) bands, with Gaussian fitting means calculated. Different waveforms during seizures were categorized into eight types using PCA and k-means clustering, with waveform and RMS heatmaps plotted. Three categories highly correlated with seizures were analyzed, producing delay heatmaps and spatial flow diagrams for the affected regions. Delay heatmaps were created by calculating the time differences between peak occurrences, illustrating the propagation of seizure waveforms. Spatial flow diagrams were generated by calculating the gradient of delay heatmaps, depicting the direction of seizure wave propagation.

## Acknowledgements

This work is supported by the Beijing Municipal Natural Science Foundation (L246017, to G.Z. and X.S.; Z220015, to L.Y.), the National Natural Science Foundation of China (NSFC) (T2425003 and 52272277, to X.S.; 92164202, to M.Z.; 62304264, to H.W.; T2122010 and 52171239 to L.Y.), NeuCyber NeuroTech (Beijing) Company (NC-2023-HE-02, to X.S.), Tsinghua University Initiative Scientific Research Program (2024Z02ORD001, to X.S.), the Science and Technology Program of Guangzhou City (2024B01J0079, to M.X. and L.Z.), the Beijing Nova Program (20230484254, to H.D.). The device fabrication and characterization work is supported by Center of Intelligent Sensing and Precise Testing Technology in Optics and Photonics, Beijing Institute of Technology and Tsinghua Nanofabrication Technology Center.

## Author contributions

Y. X., J. G., M. L., M. X., L. Z., and X. S. performed device design, fabrication and characterization. Y. X., M. L., J. G., L. Y., X. Z., Y. G, J. L, Y. C., X. L., M. Z. and X. S. designed and tested circuits. Y. X., Z. Z., M. L., J. G., Z. H., X. F., Q. L., Y. W. and X. S. designed and performed biological experiments. Y. X., M. L., J. G, L. M. and X. S. performed data processing. L.Z., X. L., M. Z., H. W., H. D., L. Y., G. Z. and X. S. provided tools and supervised the research. Y. X. and X. S. wrote the paper in consultation with other authors.

## Data and materials availability

All data needed to evaluate the conclusions in the paper are present in the paper and/or the Supplementary information

---

**Figure S1.**
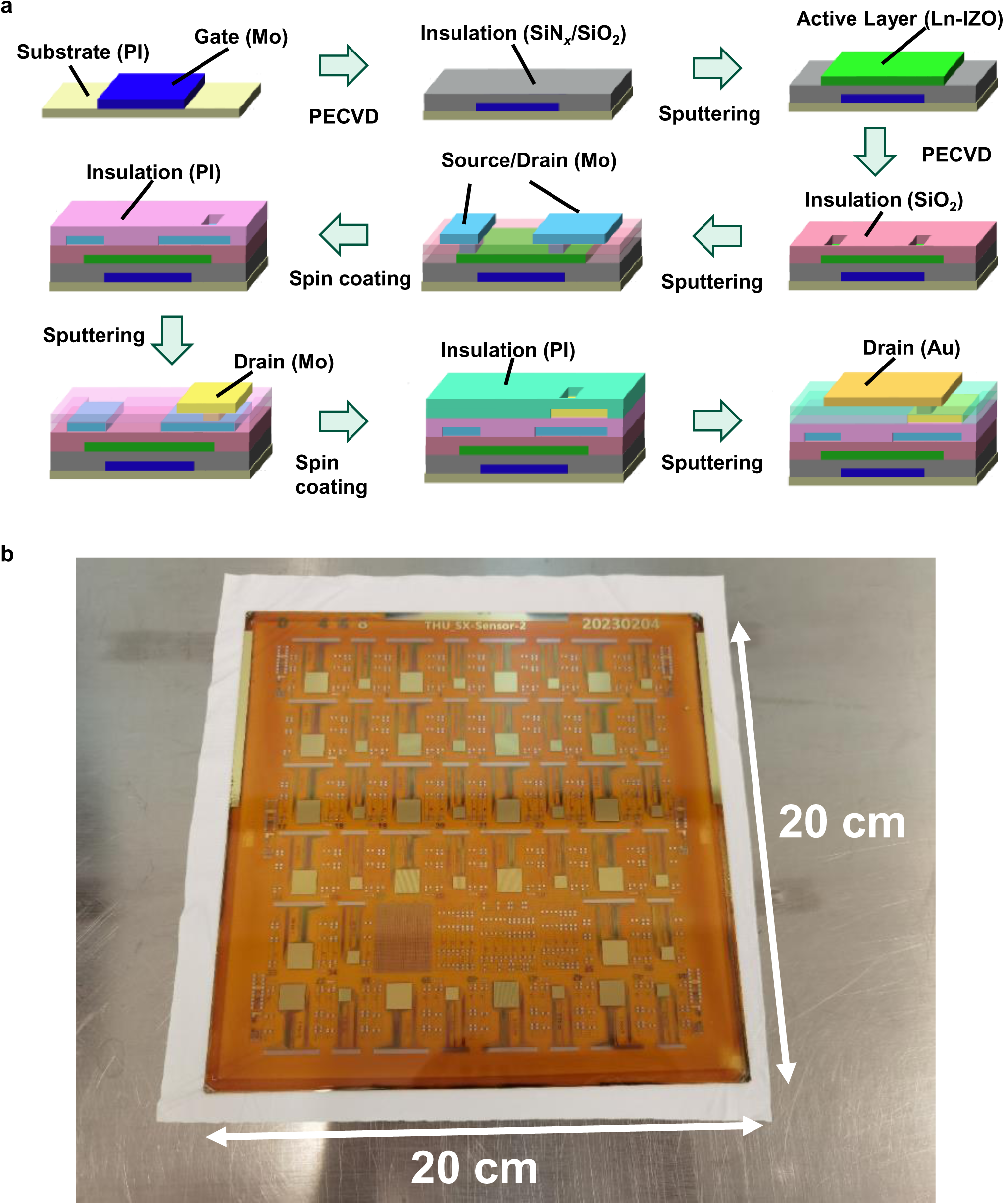
**a.** Process flow for device fabrication. **b.** Photograph of a fully-formed, large-area (20 cm × 20 cm) flexible sheet comprising multiple (>40) TFT arrays of different geometries laminated on glass.

**Figure S2.**
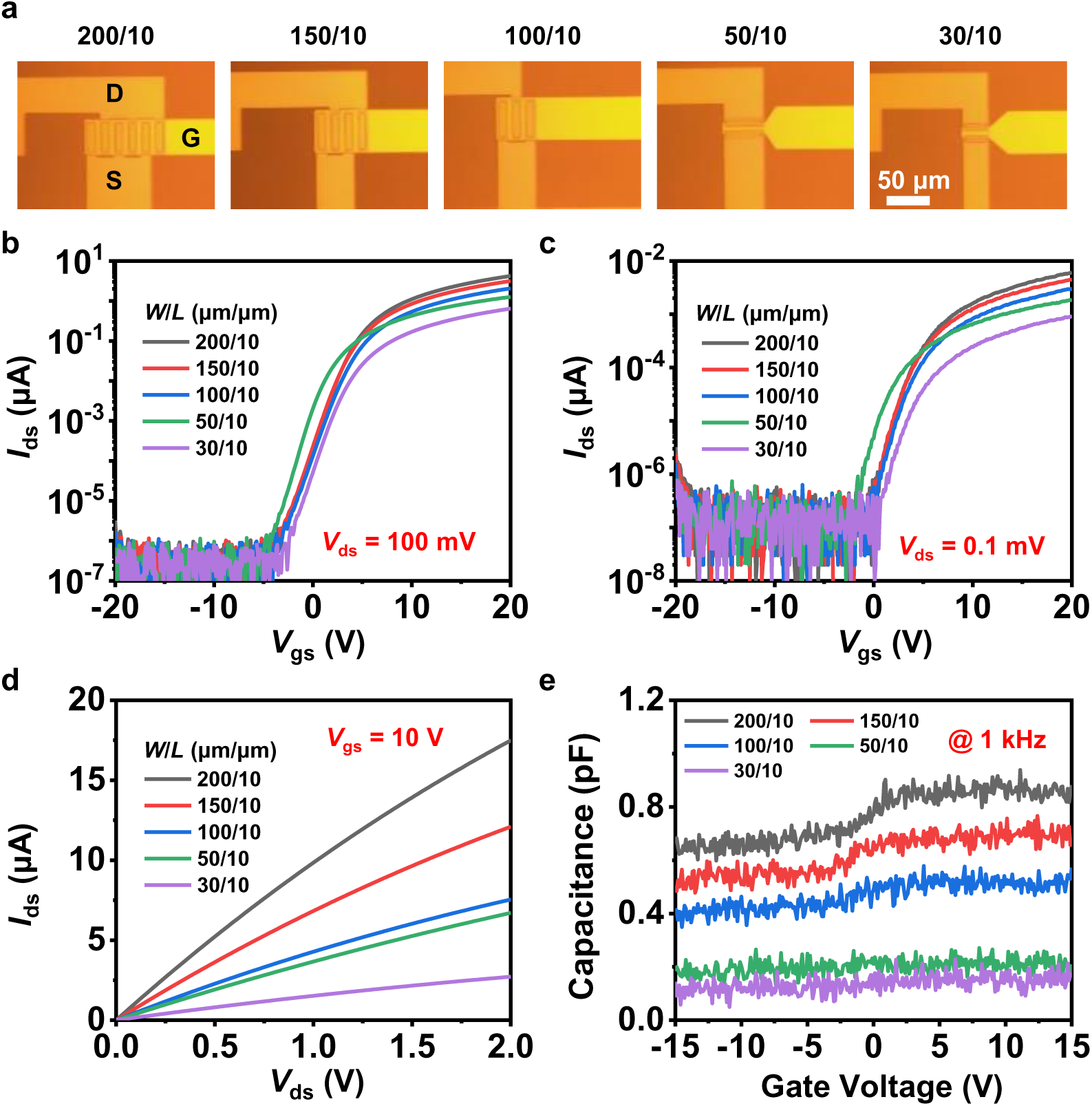
Photographs and electrical characteristics of Ln-IZO TFTs with different channel width-to-length ratios (*W*/*L*). **a.** Microscopic images (top-view) of TFTs with different geometries *W*/*L* (μm/μm). G, D, S represent gate, drain and source electrodes, respectively. **b, c.** Transfer (*I*_ds_ vs. *V*_gs_) characteristic curves at (b) *V*_ds_ = 100 mV and (c) *V*_ds_ = 0.1 mV. **d.** Output (*I*_ds_ vs. *V*_ds_) characteristic curves at *V*_gs_ = 10 V. **e.** Capacitance characteristic curves measured at 1 kHz.

**Figure S3.**
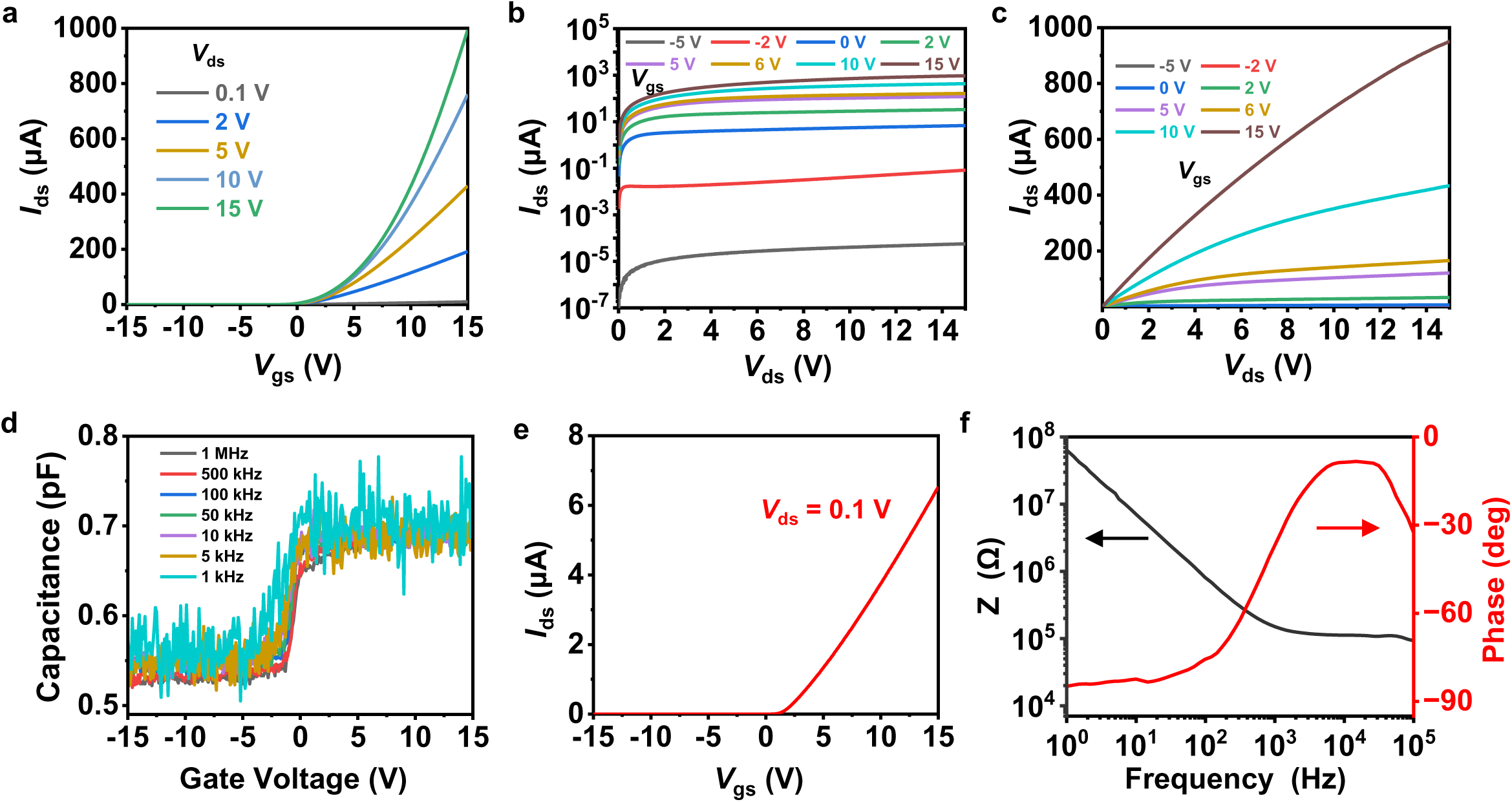
Electrical characteristics of Ln-IZO TFTs with a channel width-to-length ratio *W*/*L* = 200 μm/10 μm. **a.** Transfer (*I*_ds_ vs. *V*_gs_) characteristic curves with various *V*_ds_. **b, c.** Output (*I*_ds_ vs. *V*_ds_) characteristic curves with various *V*_gs_, plotted on (b) logarithmic and (c) linear scales. **d.** Capacitance characteristic curves at various frequencies. **e.** Transfer (*I*_ds_ vs. *V*_gs_) characteristic curve at *V*_ds_ = 0.1 V. **f.** Impedance (vs. Frequency) characteristic curve at *V*_gs_ = 6 V, measured by immersing a TFT into the PBS solution.

**Figure S4.**
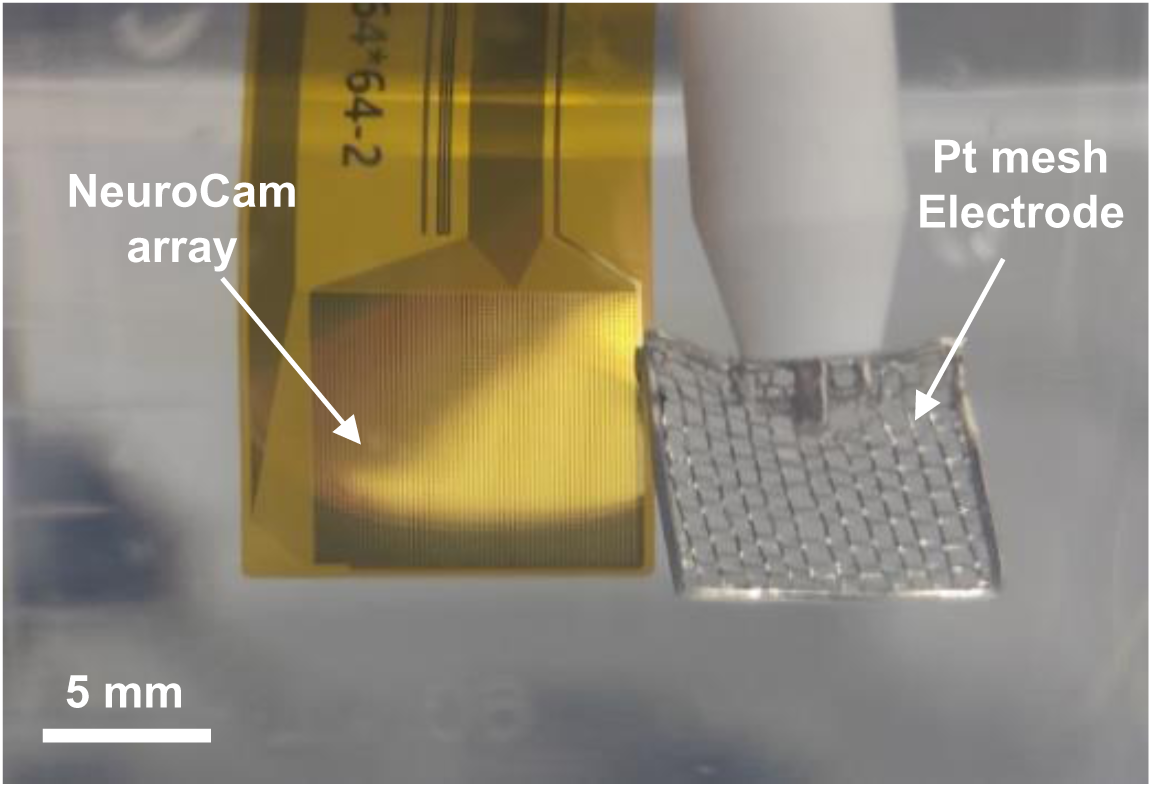
Photograph of the device fully immersed in PBS during the soak test.

**Figure S5.**
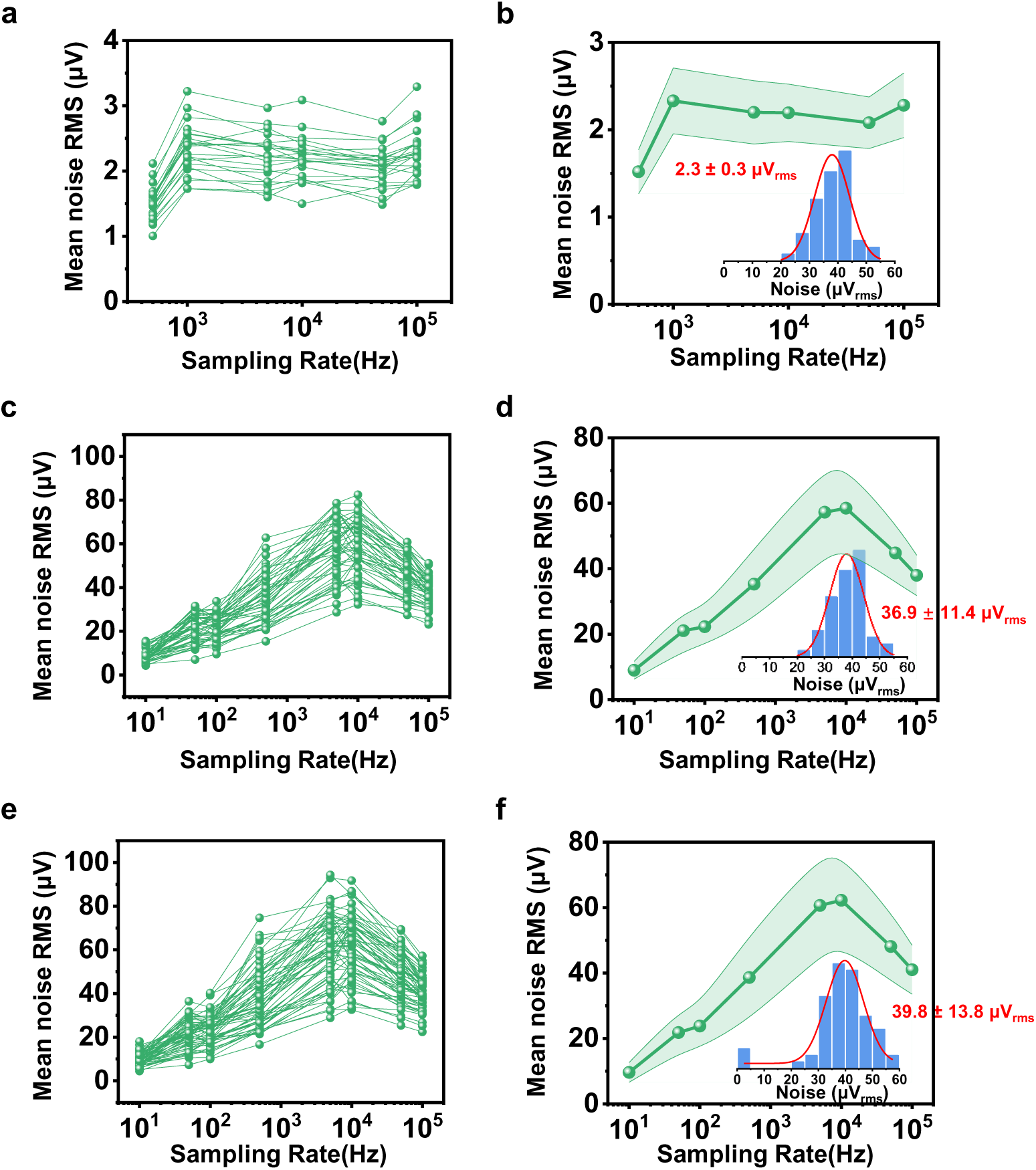
Noise characteristics of the NeuroCam system at the static mode. The array is immersed in PBS, and the input signal is applied using a platinum mesh electrode. One G line is kept on. **a.** Mean noise RMS result over 1 second from 23 S lines at various sampling rate. **b.** Statistical results of mean noise RMS for these 23 channels. Inset: Histogram and Gaussian fit curve of noise levels for all channels at a sample rate of 100 kS/s. **c.** Mean noise RMS result over 1 second from all 48 S lines at various sampling rate. **d.** Statistical results of mean noise RMS for these 48 channels. Inset: Histogram and Gaussian fit curve of noise levels for all channels at a sample rate of 100 kS/s. **e.** Mean noise RMS result over 1 second from all 64 S lines at various sampling rate. **f.** Statistical results of mean noise RMS for these 64 channels. Inset: Histogram and Gaussian fit curve of noise levels for all channels at a sample rate of 100 kS/s.

**Figure S6.**
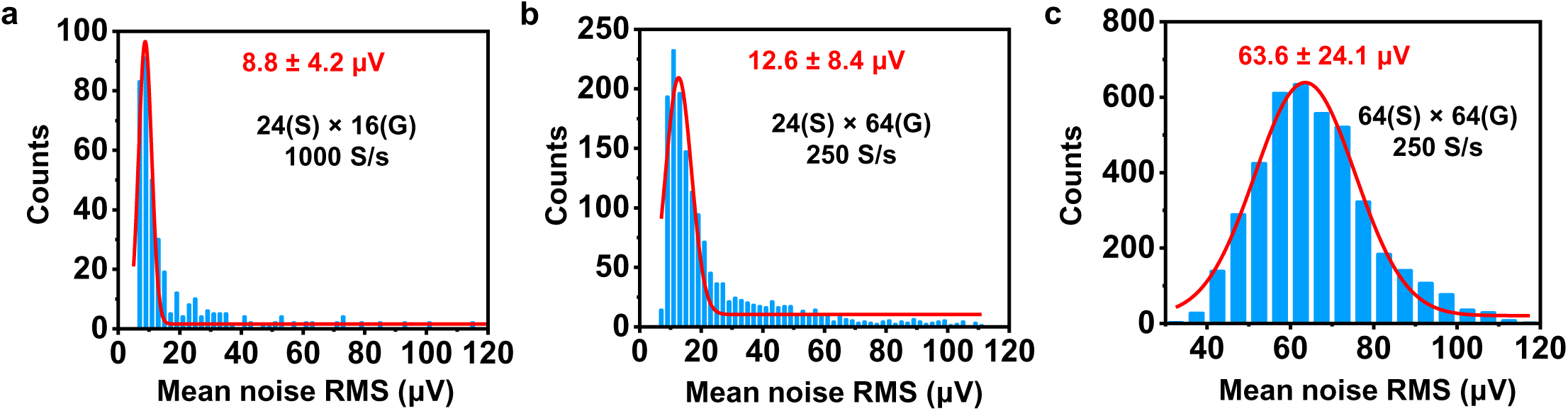
Noise characteristics of the NeuroCam system at the dynamic mode. The array is immersed in PBS, and the input signal is applied using a platinum mesh electrode. The sample rate for each S line is fixed at 100 kS/s. By adjusting the number of operated active lines (S and G), effective sample rates for each channel can be varied and corresponding noise levels are different. Effective sample rates for operating 24(S) × 16(G), 24(S) × 64(G), 64(S) × 64(G) are 1000 S/s, 250 S/s and 250 S/s, respectively.

**Figure S7.**
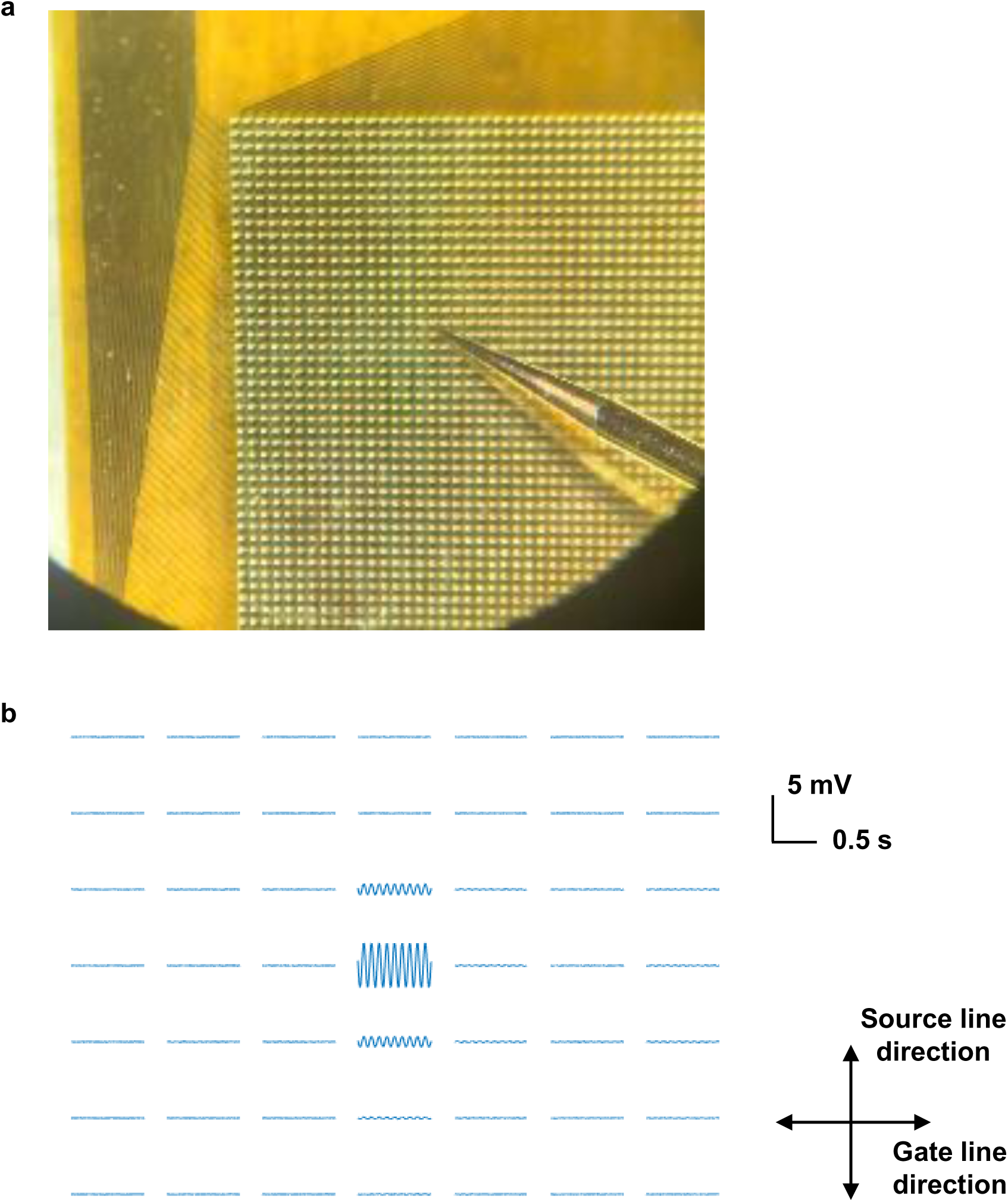
Crosstalk analysis of the NeuroCam array. **a.** Photograph showing a device array exposed in air, with a contact probe positioned on a single pixel and applying a sinusoidal signal (5 mVpp, 12 Hz). **b.** Measured output voltages from the probed pixel and its adjacent pixels. With only one point as the input signal, it can be observed that the other points are almost unaffected. Adjacent pixels in the direction of source line receives higher crosstalk signals (−6.3 dB) comparing to those in the direction of gate line (−18.6 dB), while second nearest neighboring and further pixels show minimal crosstalk signals (−22.5 dB).

**Figure S8.**
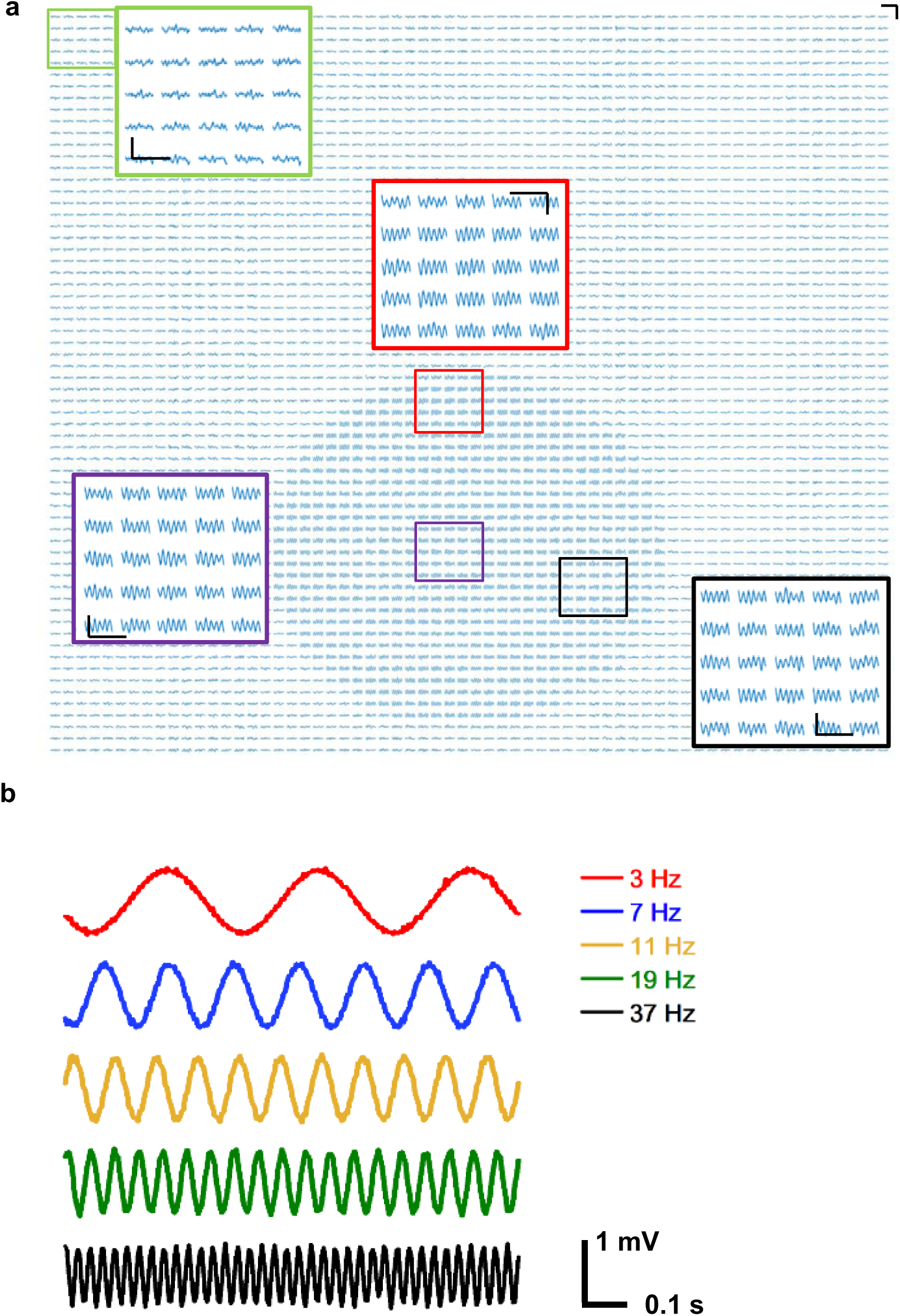
In vitro recording signals from all 4096 channels in the TFT array. **a.** A droplet of PBS is placed on the device surface, and a 1 mVpp sine signal (18 Hz) is applied into PBS. Insets show enlarged plots from different regions. Scalebars are 0.5 s and 1 mV. **b.** Signals recorded when injecting signals at different frequencies (3 Hz, 7 Hz, 11 Hz, 19 Hz, and 37 Hz) in PBS solution environment, corresponding to delta, theta, alpha, beta, and gamma bands of ECoG signals, respectively.

**Figure S9.**
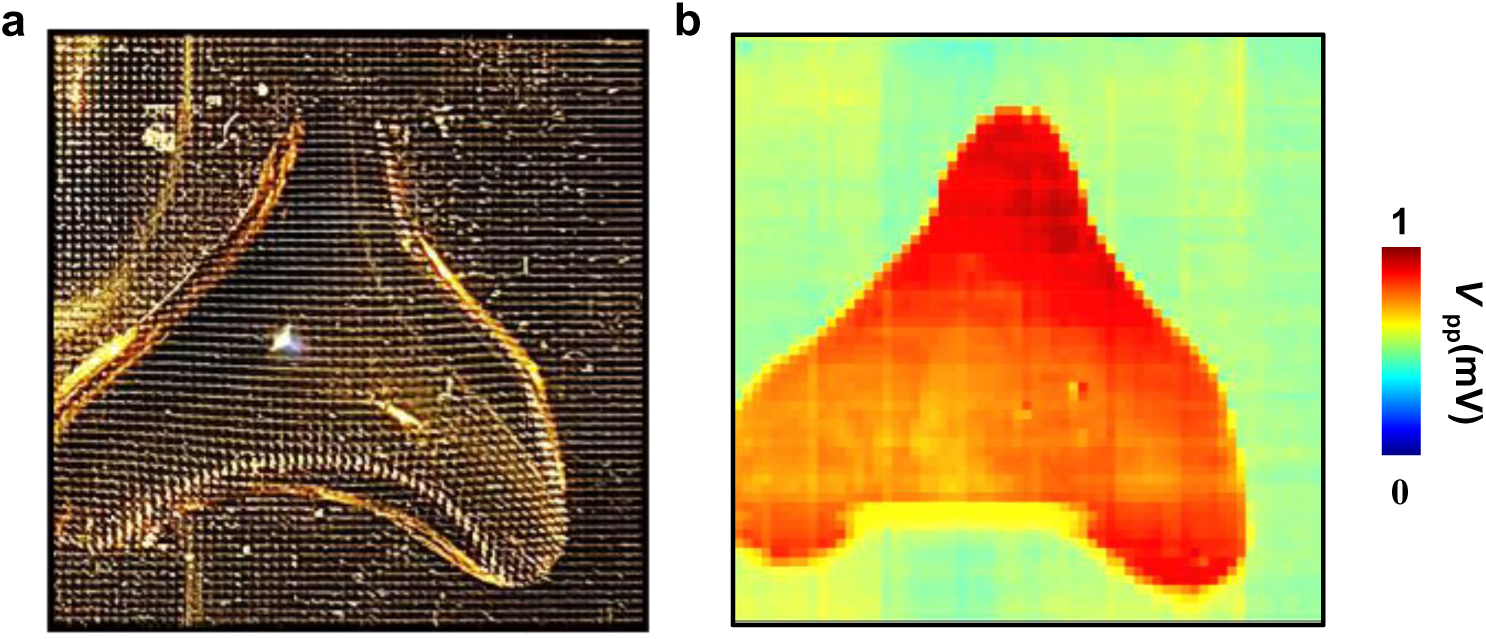
**a.** Photograph of a NeuroCam array with a droplet with a complex shape of PBS on its surface. **b.** Mapping results of spatially resolved output signals when injecting a 1 mVpp sine signal (12 Hz) into PBS.

**Figure S10.**
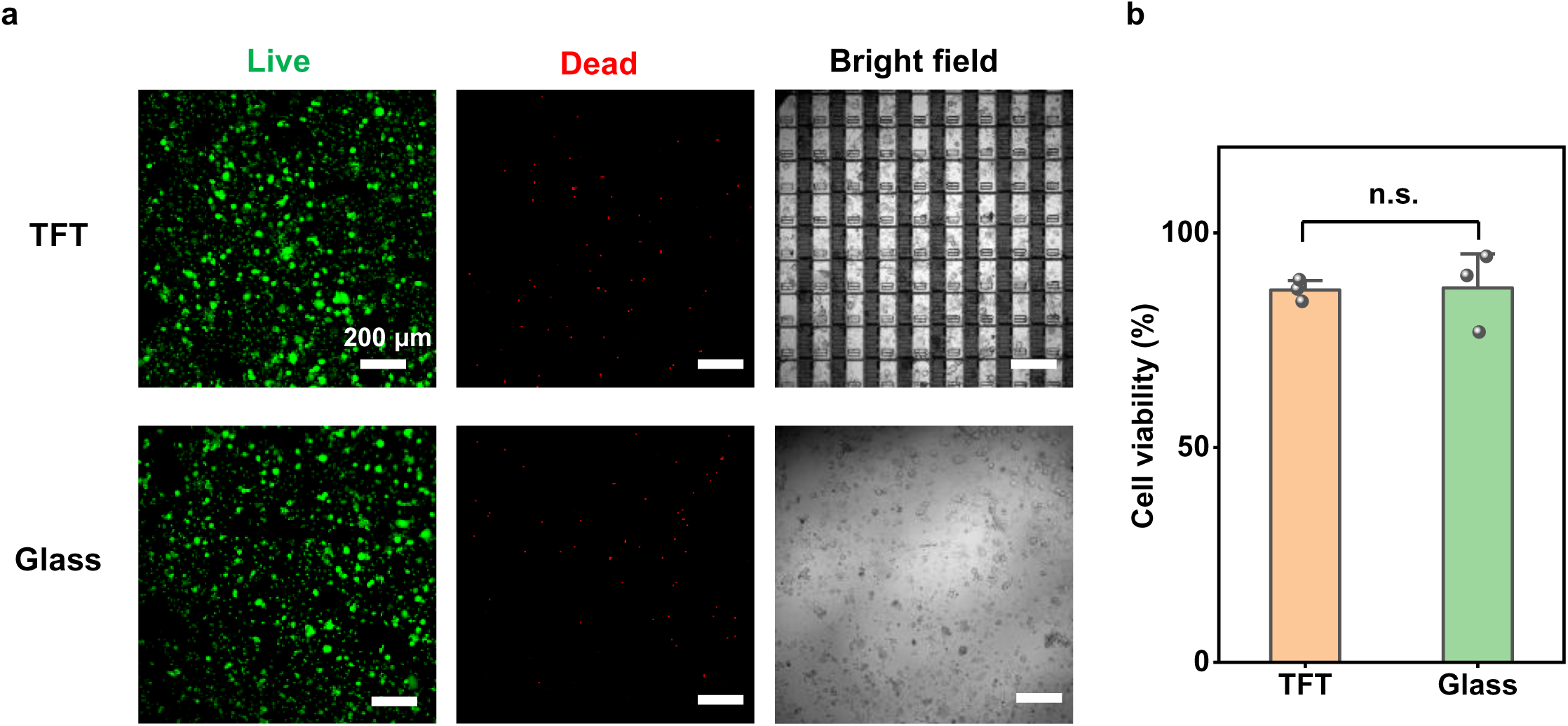
Biocompatibility of the TFT array. **a.** Live/dead fluorescence assay performed on DRG cells cultured on the TFT and glass for 24 h. Green (Calcien AM) and red (Propidium Iodide, PI) denote live cells and dead cells, respectively. **b.** Averaged viability (%) of DRG cells cultured on the TFT and glass. Data are presented as mean ± s.e.m. (*n* = 3 devices) and analyzed by unpaired *t*-test. n.s., not significant.

**Figure S11.**
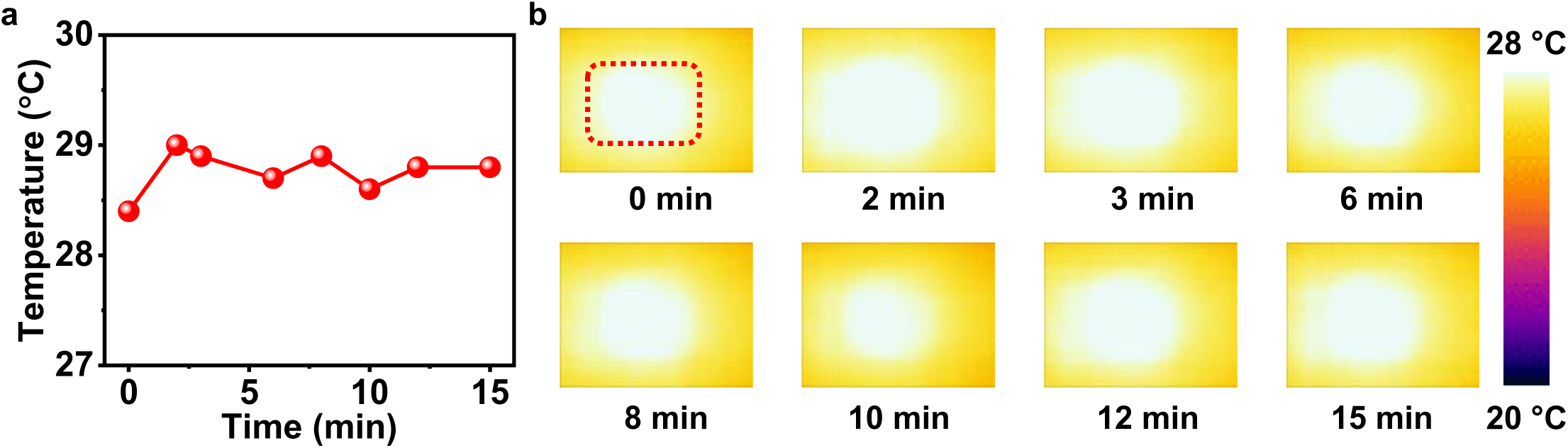
Thermal properties of the TFT array during operation. **a.** Plot of measured maximum surface temperature as a function of operation time. **b.** Thermography of a device array at different time points. During device operation, the maximum temperature is less than 29 °C.

**Figure S12.**
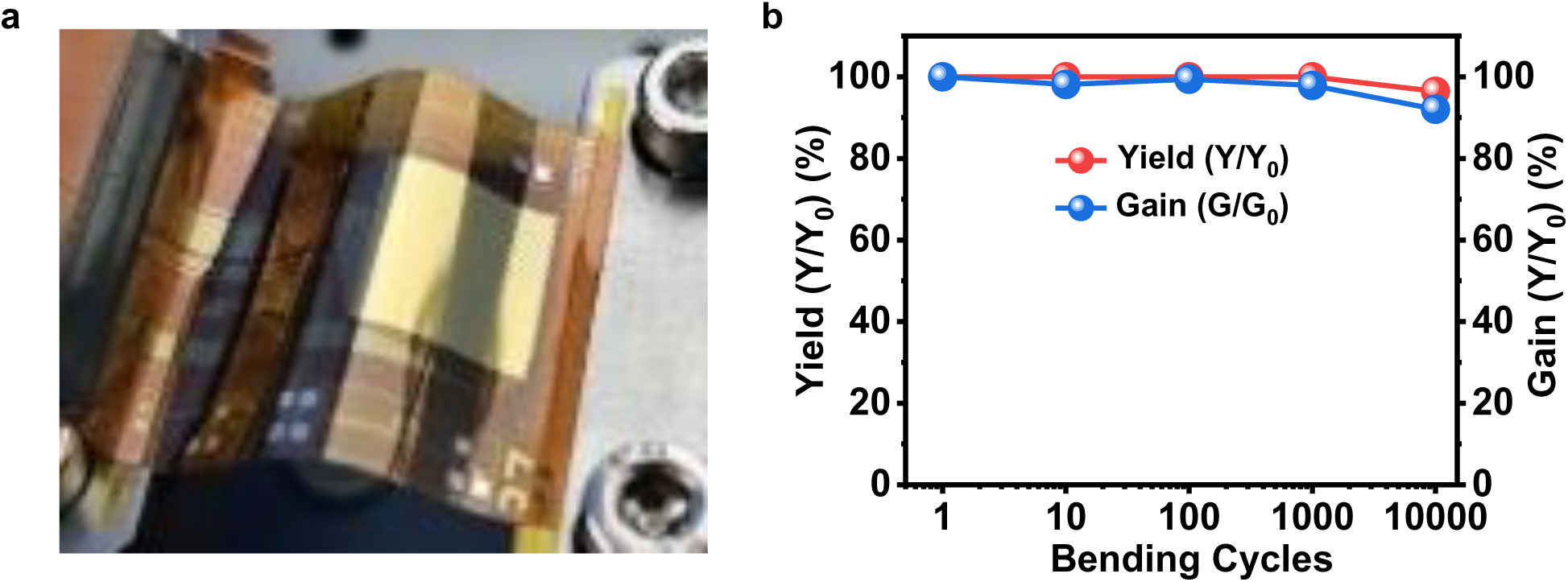
Mechanical properties of the TFT array. **a.** Experimental setup for dynamic bending test. **b.** Yield and average gain variation of the array under different bending cycles. The minimum bending radius is 5 mm. Yield is defined as the ratio of the number of channels functioning properly after bending to the number of channels functioning before bending. Gain is defined as the ratio of the average signal amplitude of all functioning channels after bending to the average signal amplitude before bending.

**Figure S13.**
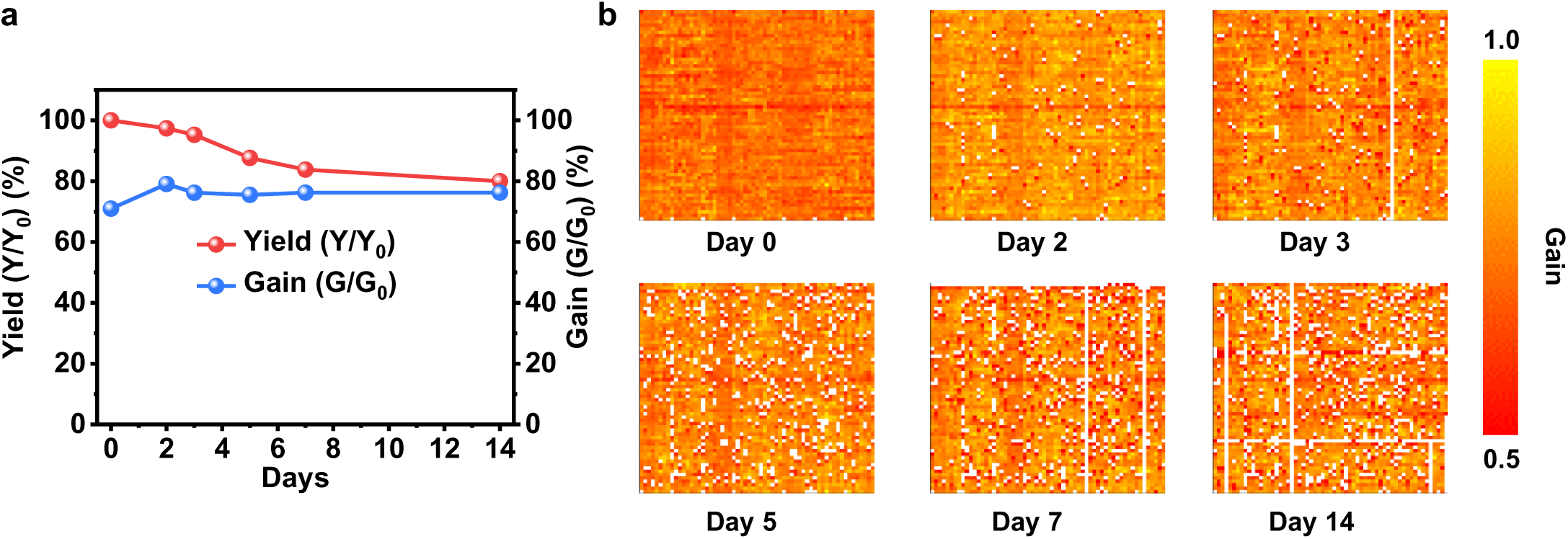
Performance of the TFT array during chronic soak test in PBS solution at 37 °C. **a.** Variation of device yield and gain over soaking time. **b.** Measured spatial distribution of gain for all channels at different time points.

**Figure S14.**
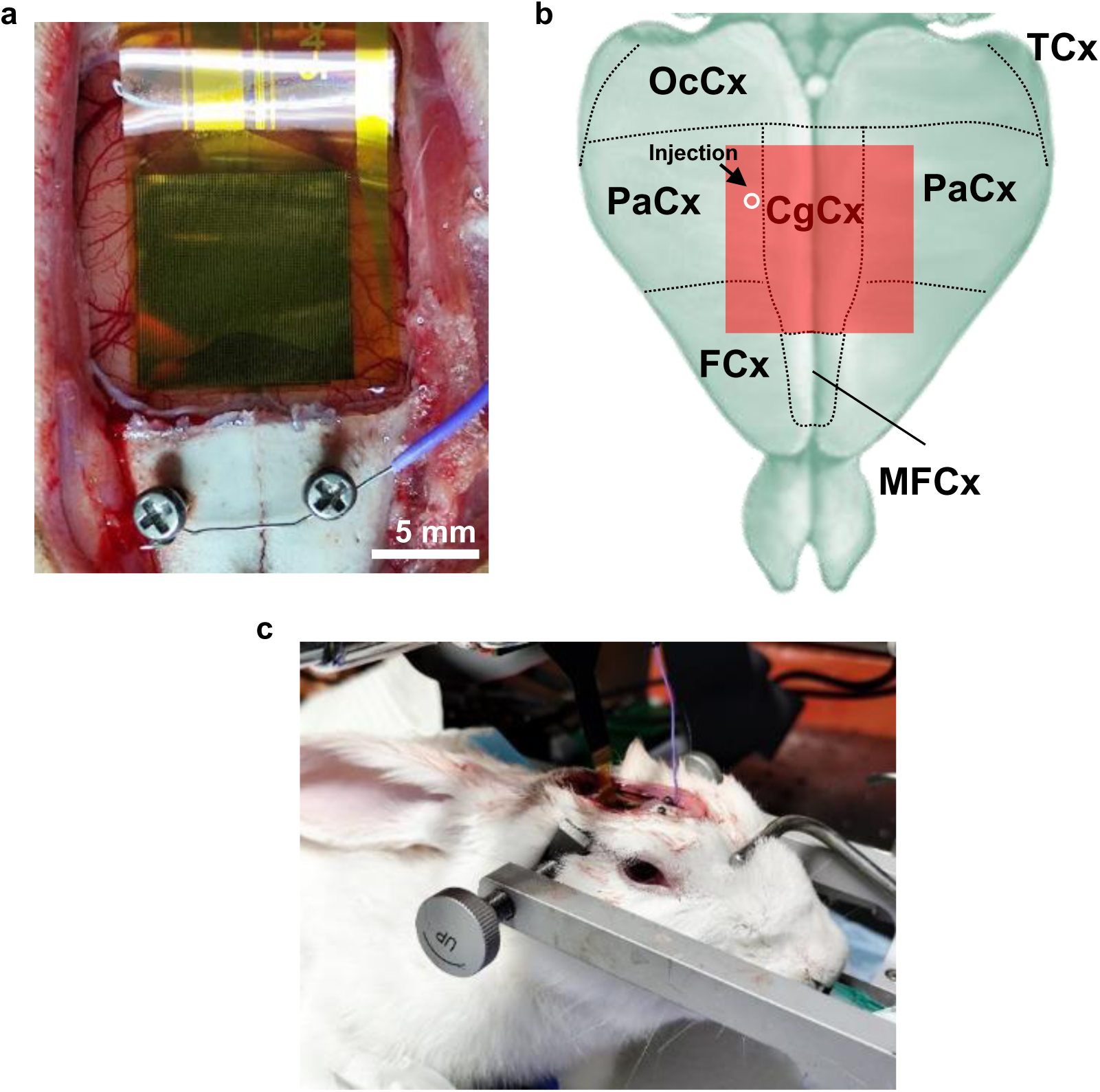
Setup for in vivo experiment using a rabbit model. **a.** A NeuroCam array placed on the cortex. **b.** Schematic illustration of the device array overlaid with segmented cortical areas of the rabbit brain (dorsal view), including temporal cortex (TCx), occipital cortex (OcCx), parietal cortex (PaCx), cingulate cortex (CgCx), frontal cortex (FCx), medial frontal cortex (MFCx). The arrow indicates the location of penicillin injection. **c.** Wired device array used in an anesthetized rabbit. Reference: E. Munoz-Moreno *et al.*, A Magnetic Resonance Image Based Atlas of the Rabbit Brain for Automatic Parcellation. *PLoS ONE* **8**, e67418 (2013).

**Figure S15.**
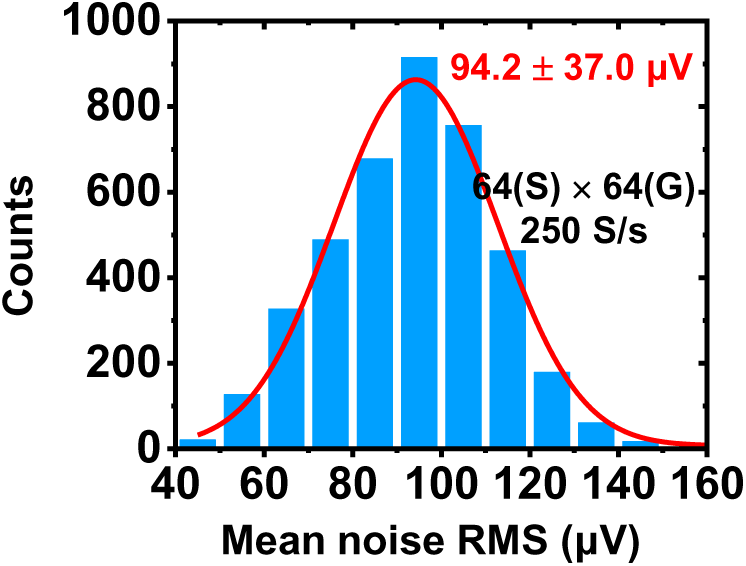
Noise characteristics of the NeuroCam system measured on the rabbit cortex. The 64 × 64 active channels are sampled at a rate of 250 S/s. It should be noted that these signals encompass the noise associated with the device array and the recording system, as well as the animal’s ECoG activity at the rest state.

**Figure S16.**
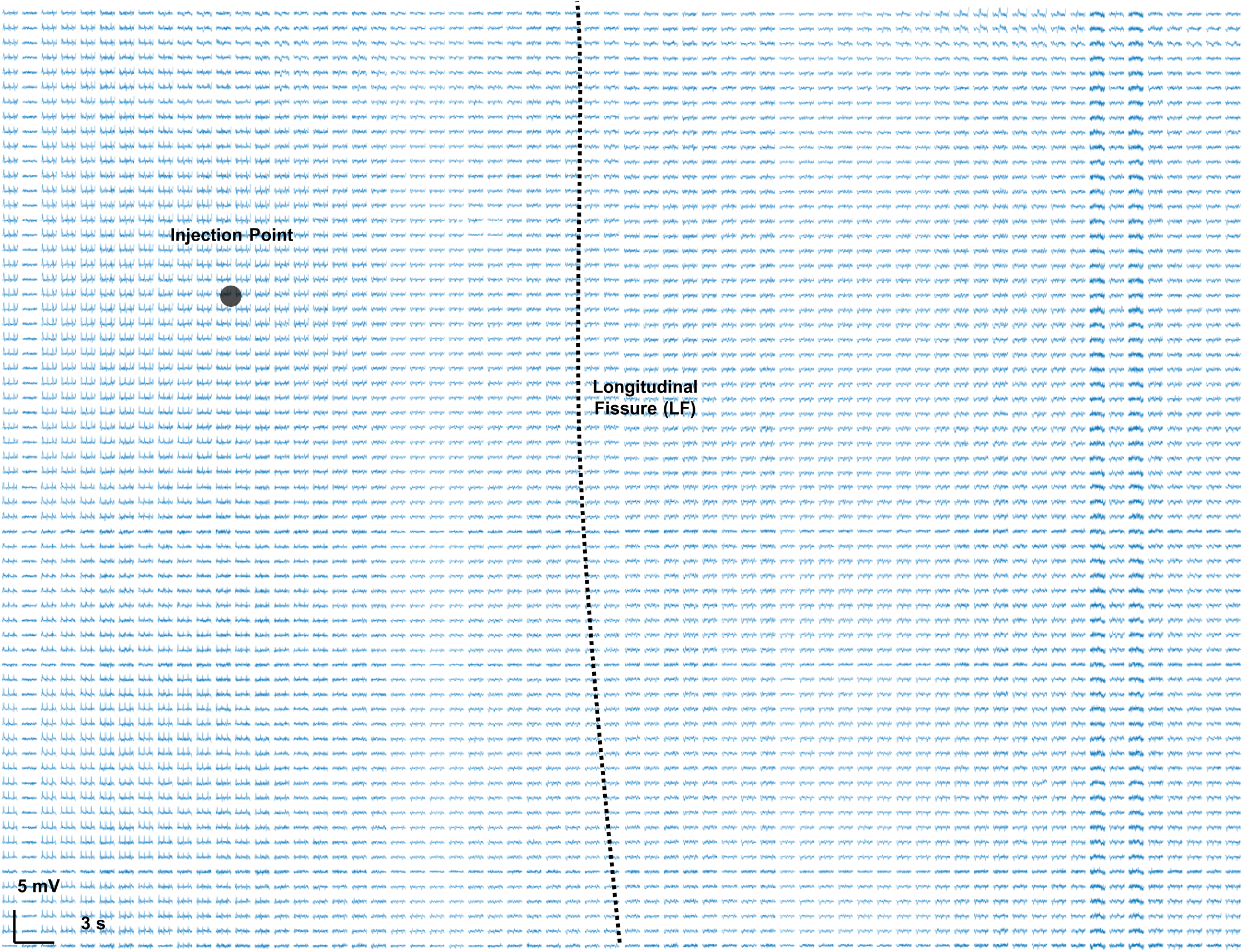
Representative ECoG activities recorded from all 4096 channels in the array, during status epilepticus in the rabbit.

**Figure S17.**
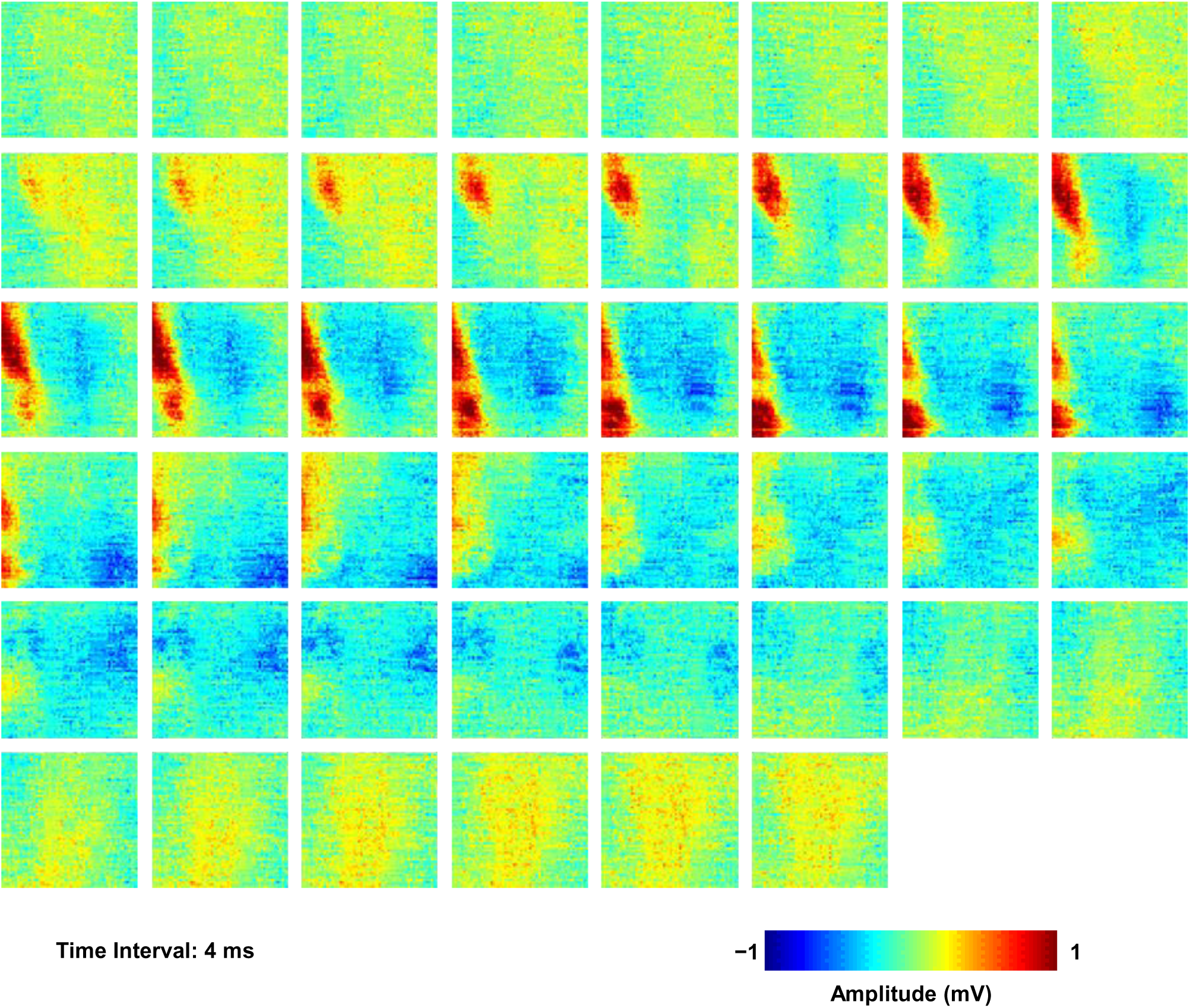
Time-lapse mapping results of a representative epileptic spike within 180 ms.

**Figure S18.**
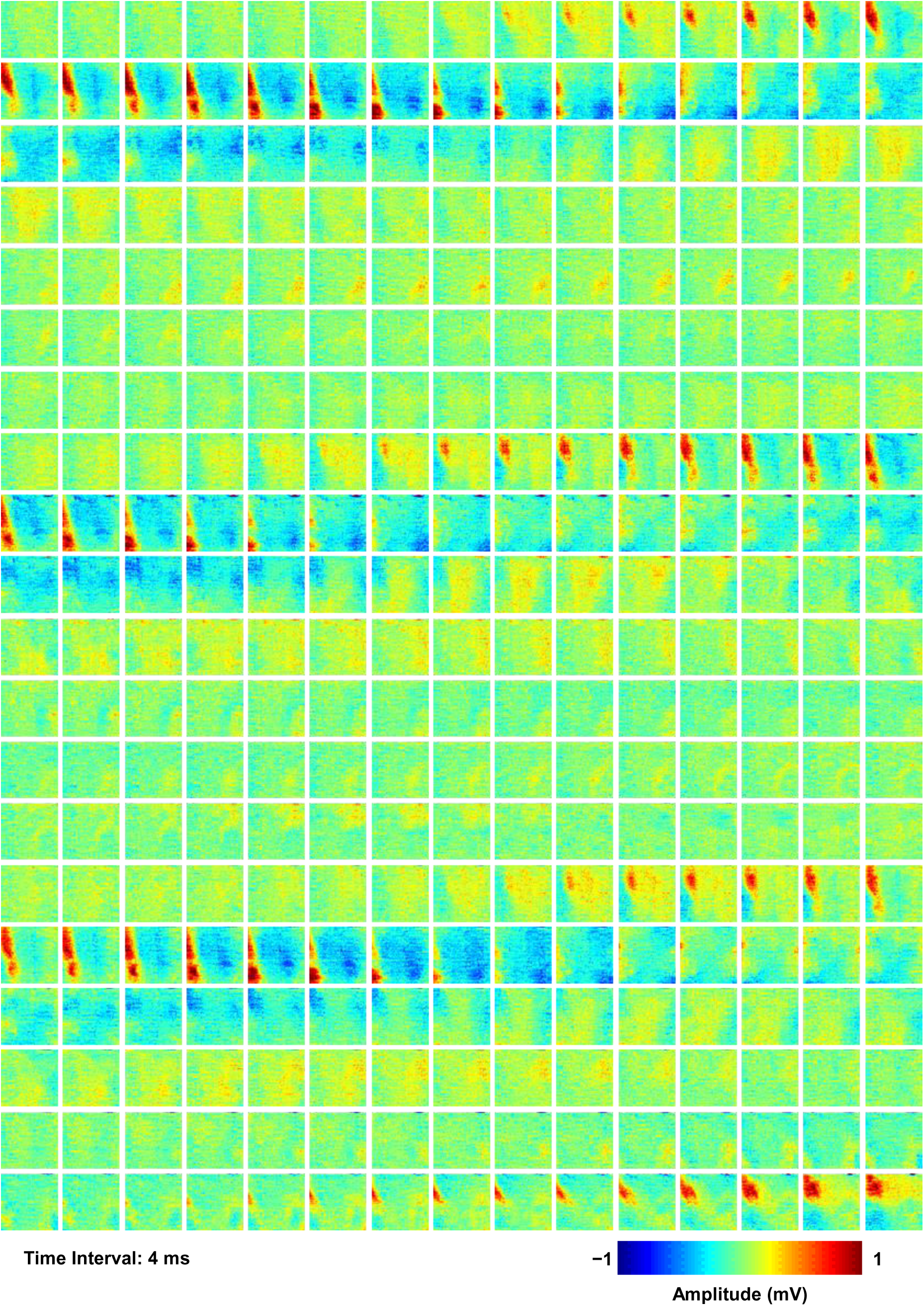
Time-lapse mapping results of three continuous epileptic spike waves within 1.2 s.

**Figure S19.**
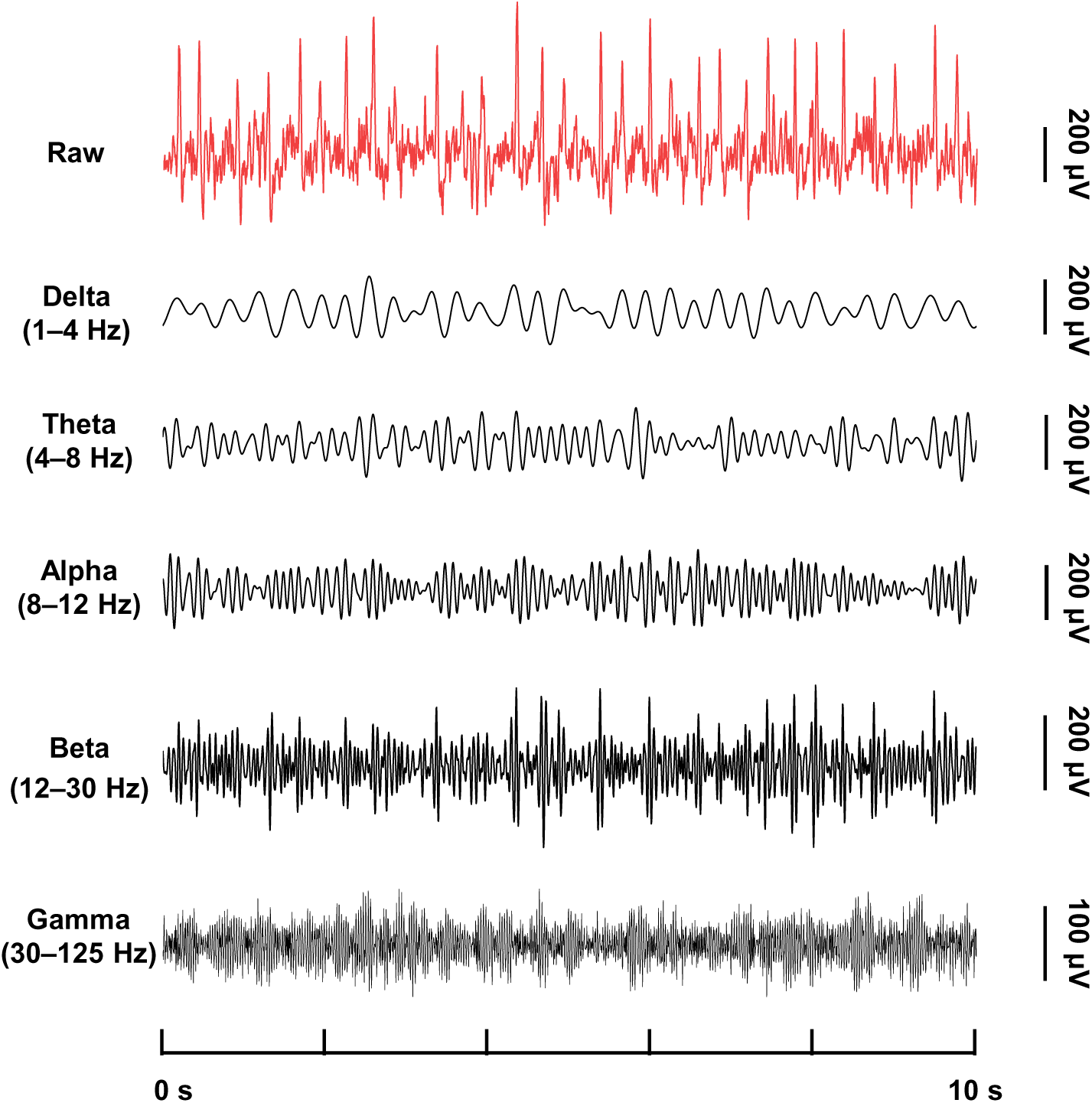
Raw ECoG data registered by a representative channel during epileptic seizures, and corresponding results in different frequency bands: Delta (1–4 Hz), Theta (4–8 Hz), Alpha (8–12 Hz), Beta (12–30 Hz), and Gamma (30–125 Hz).

**Figure S20.**
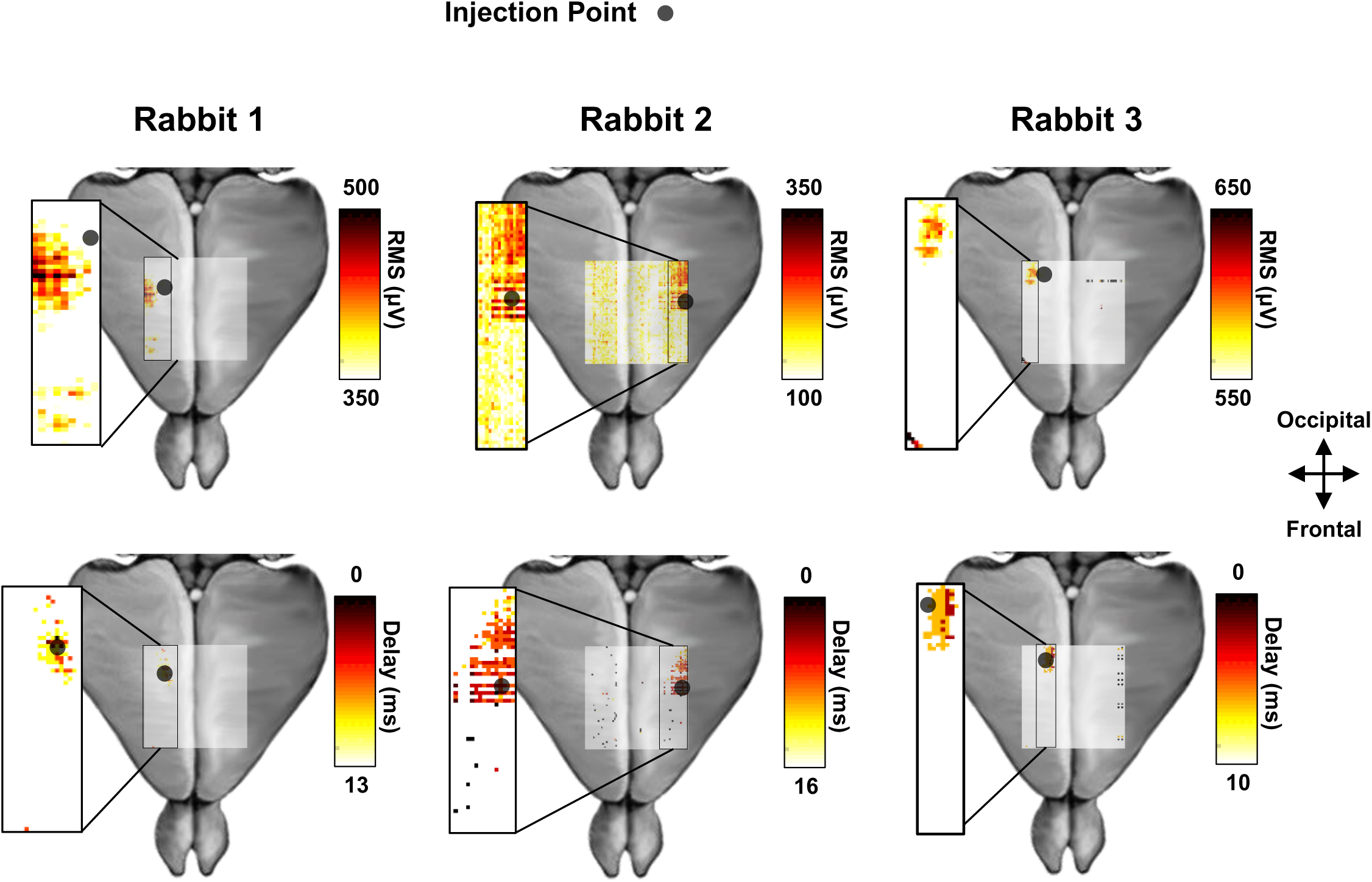
Mapping of ECoG signal amplitudes and delays for all the channels during epileptic spikes in three rabbits with penicillin injected in different locations. **Top:** RMS signals from the most active channels in the array and their spatial locations. **Bottom:** Delay of signals indicating channels with the earliest bursting in the array and their spatial locations.

**Figure S21.**
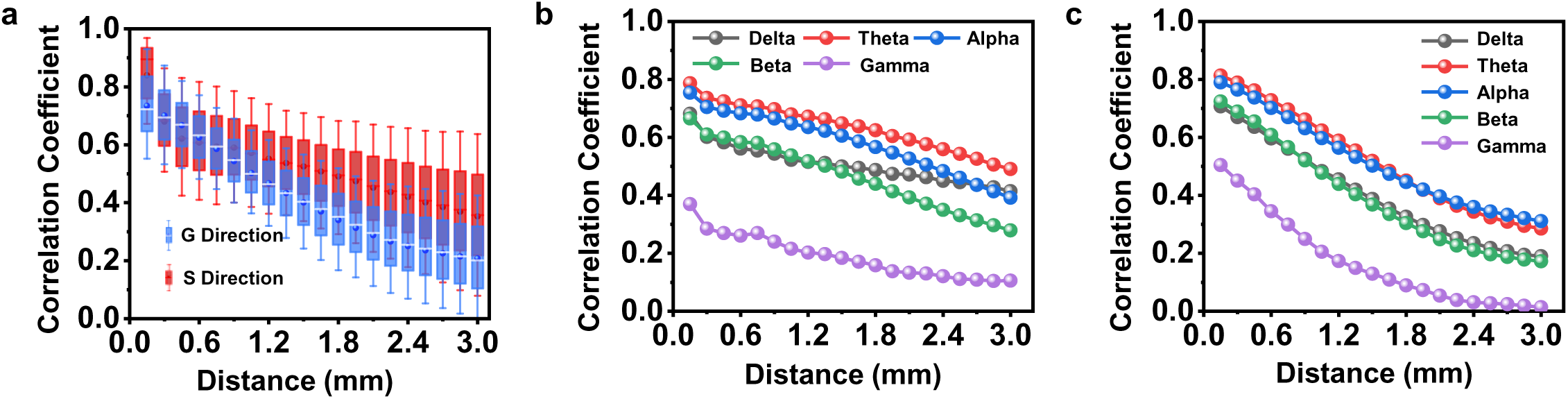
Spatial correlation of signals registered by different channels in the NeuroCam array. **a.** Relationship between correlation coefficients and inter-electrode distances, in the G and S directions. **b-c.** Relationship between correlation coefficients and inter-electrode distances, in different frequency bands. **b.** G direction and **c.** S direction.

**Table S1.**
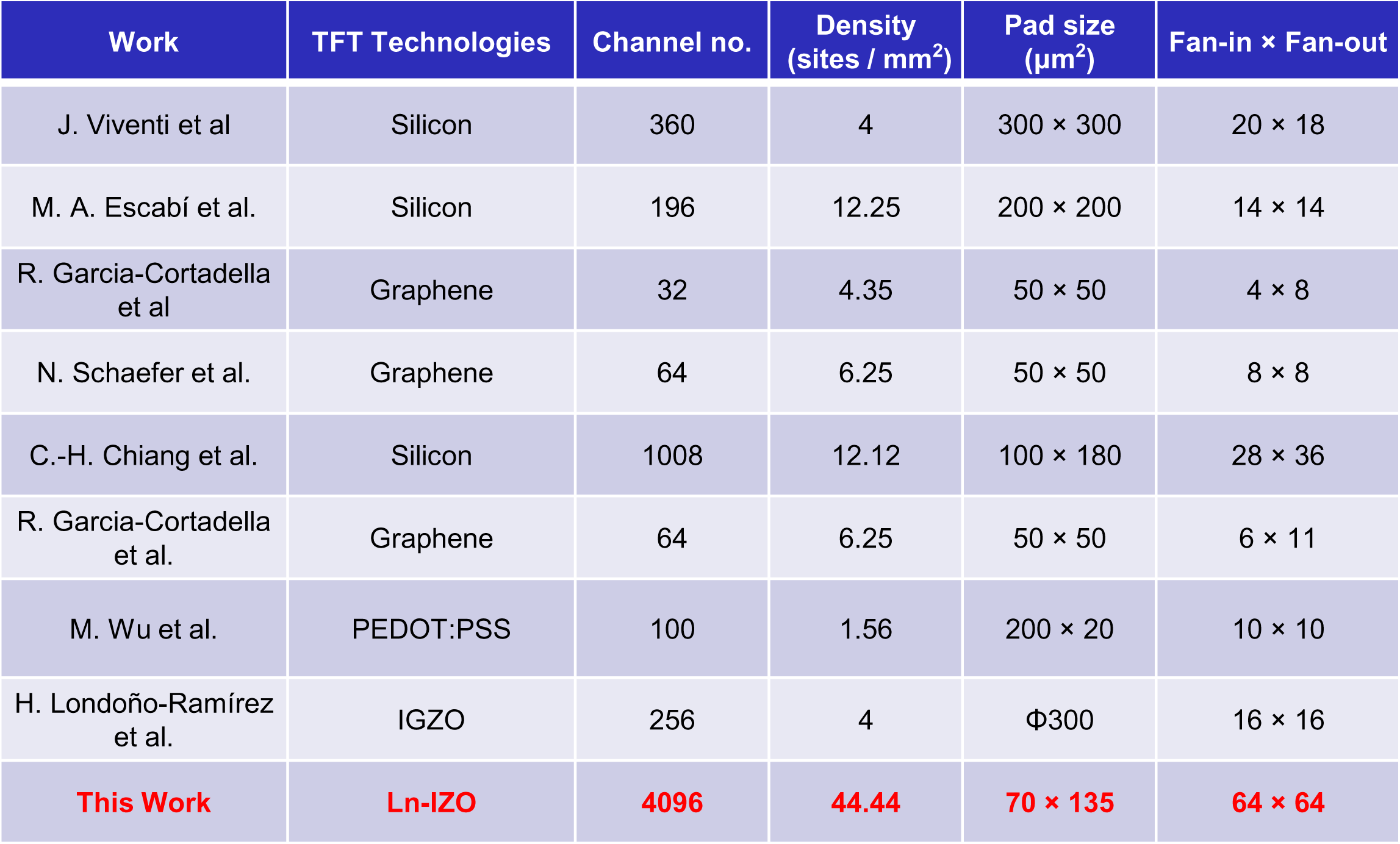
Comparison of characteristic parameters between this work and other multiplexed TFT-based μECoG arrays performed in vivo validation.

| Work | TFT Technologies | Channel no. | Density (sites / mm<sup>2</sup>) | Pad size (μm<sup>2</sup>) | Fan-in × Fan-out |
| --- | --- | --- | --- | --- | --- |
| J. Viventi et al | Silicon | 360 | 4 | 300 × 300 | 20 × 18 |
| M. A. Escabí et al. | Silicon | 196 | 12.25 | 200 × 200 | 14 × 14 |
| R. Garcia-Cortadella et al | Graphene | 32 | 4.35 | 50 × 50 | 4 × 8 |
| N. Schaefer et al. | Graphene | 64 | 6.25 | 50 × 50 | 8 × 8 |
| C.-H. Chiang et al. | Silicon | 1008 | 12.12 | 100 × 180 | 28 × 36 |
| R. Garcia-Cortadella et al. | Graphene | 64 | 6.25 | 50 × 50 | 6 × 11 |
| M. Wu et al. | PEDOT:PSS | 100 | 1.56 | 200 × 20 | 10 × 10 |
| H. Londoño-Ramírez et al. | IGZO | 256 | 4 | Φ300 | 16 × 16 |
| **This Work** | **Ln-IZO** | **4096** | **44.44** | **70 × 135** | **64 × 64** |

**Movie S1.**
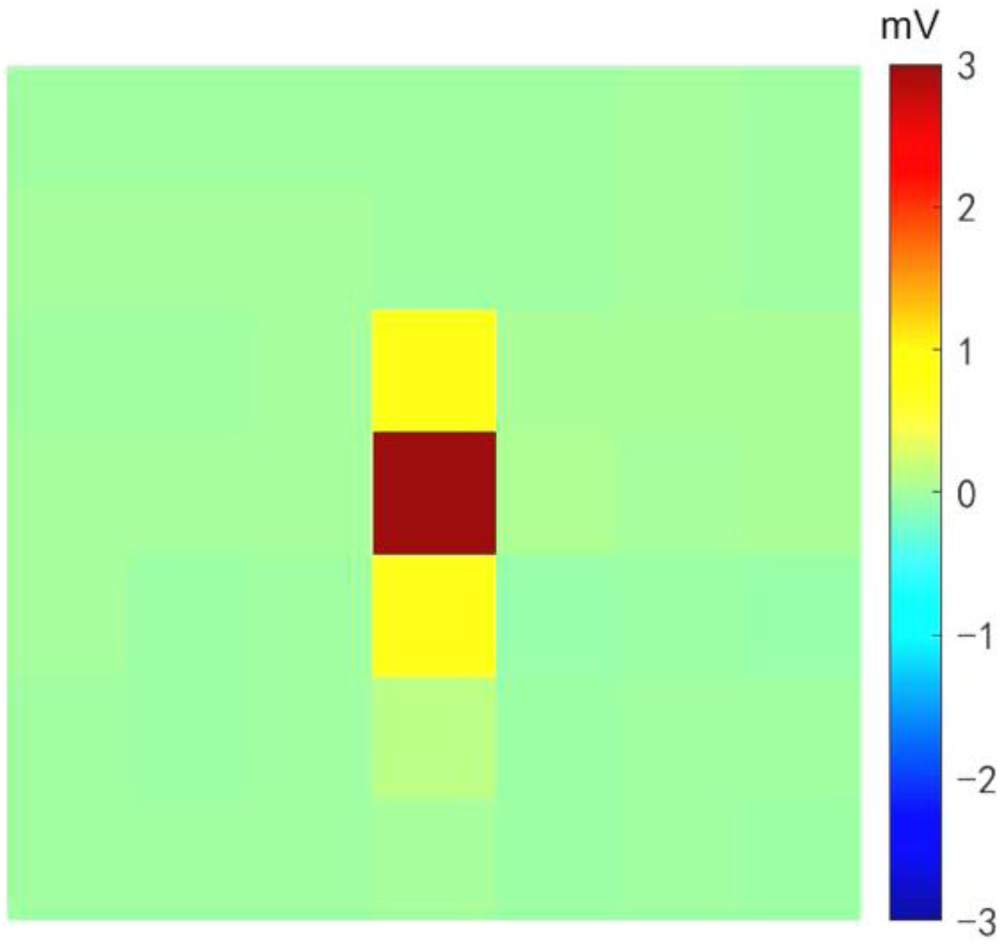
Slow motion video (0.1× speed) shows the crosstalk recorded by the NeuroCam array.

**Movie S2.**
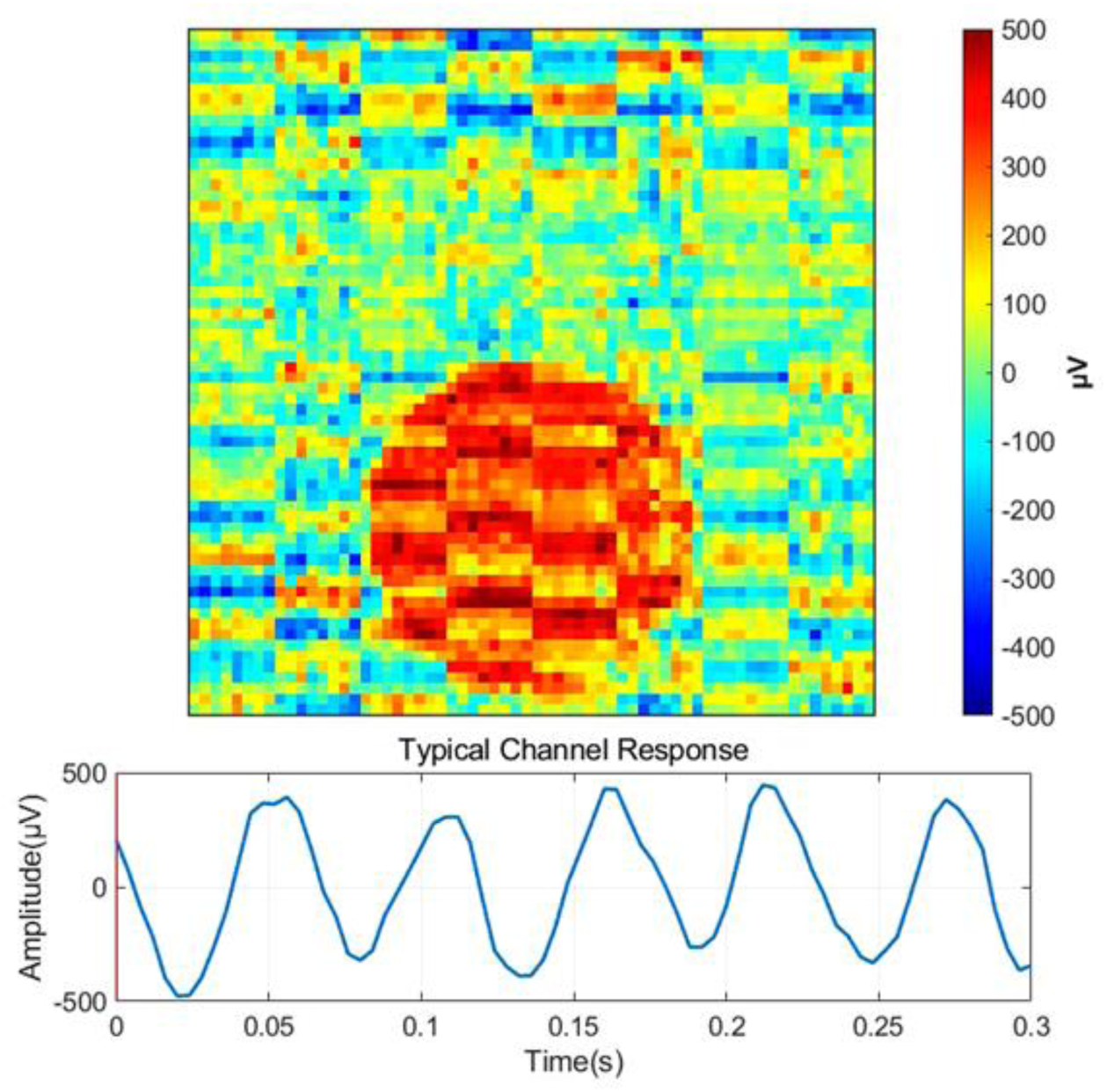
Slow motion video (0.1× speed) shows in vitro mapping results of spatially resolved output signals when injecting a 1 mVpp sine signal (12 Hz) into a PBS droplet with a circular shape on the NeuroCam array.

**Movie S3.**
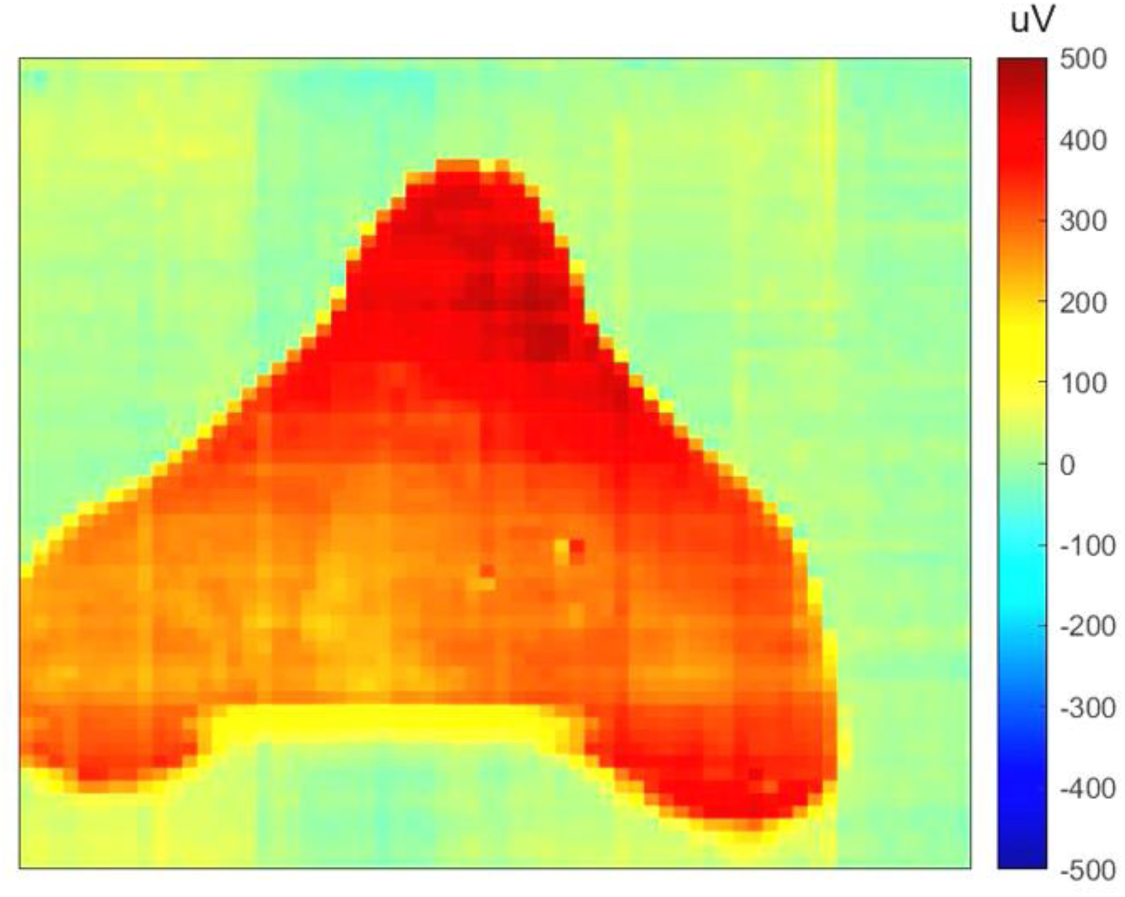
Slow motion video (0.1× speed) shows in vitro mapping results of spatially resolved output signals when injecting a 1 mVpp sine signal (12 Hz) into a PBS droplet with a more complex shape on the NeuroCam array.

**Movie S4.**
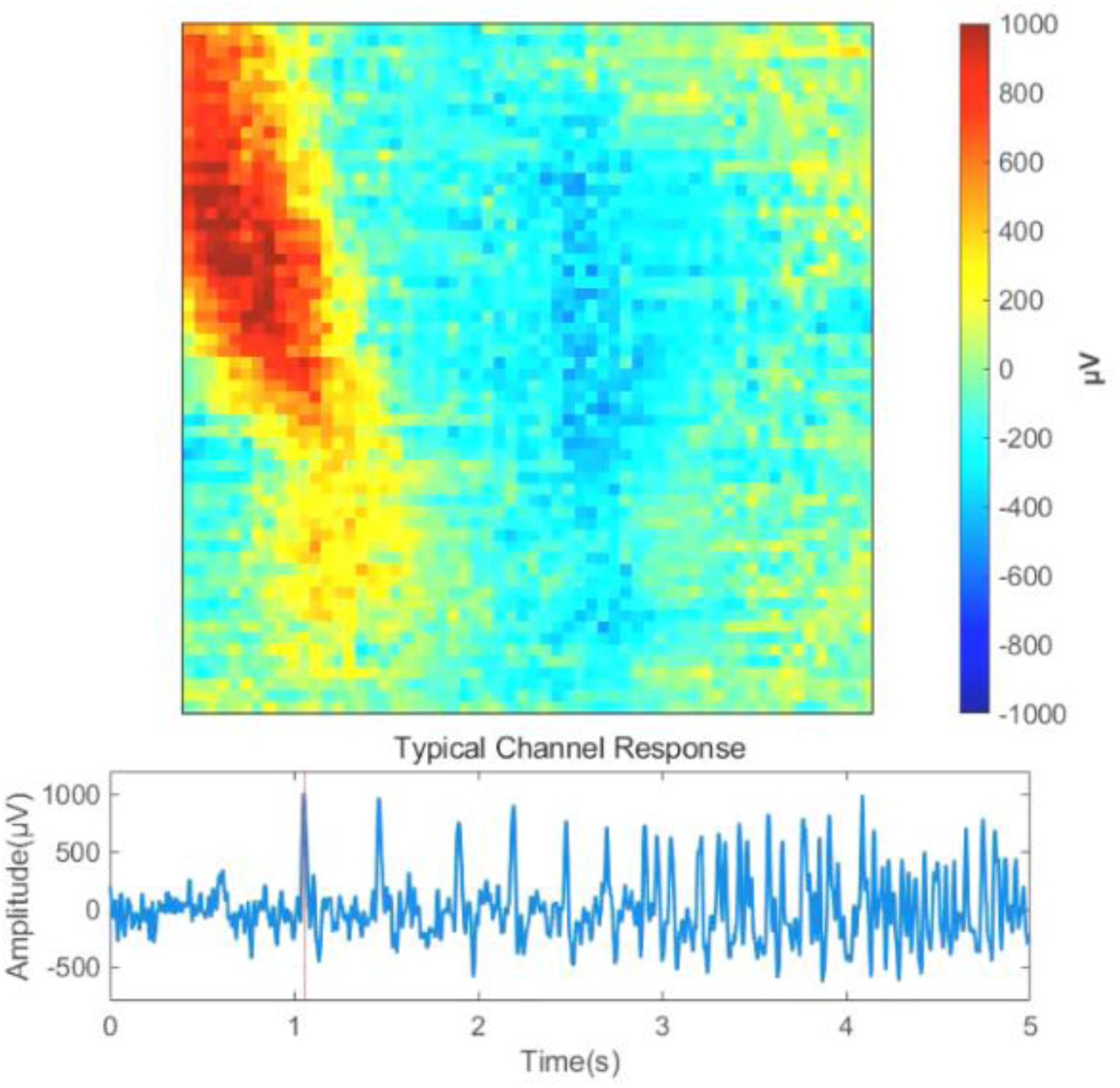
Slow motion video (0.1× speed) shows in vivo spatiotemporal evolution of epileptiform discharges captured by the NeuroCam array.

